# A nucleation-and-growth model for the packaging of genome in linear virus-like particles: impact of multiple packaging signals

**DOI:** 10.1101/2022.02.24.481677

**Authors:** René de Bruijn, P.C.M. Wielstra, Carlos Calcines-Cruz, Tom van Waveren, Armando Hernandez-Garcia, Paul van der Schoot

## Abstract

Inspired by recent experiments on the spontaneous assembly of virus-like particles from a solution containing a synthetic coat protein and double-stranded DNA, (1) we put forward a kinetic model that has as main ingredients a stochastic nucleation and a deterministic growth process. The efficiency and rate of the packaging of the DNA turn out to strongly increase by introducing proteins onto the DNA template that are modified using CRISPR-Cas techniques to bind specifically at predesignated locations, mimicking assembly signals in viruses. Our model shows that treating these proteins as nucleation-inducing diffusion barriers is sufficient to explain experimentally observed increase in encapsulation efficiency, but only if the nucleation rate is sufficiently high. We find an optimum in the encapsulation kinetics for conditions where the number of packaging signals is equal to the number of nucleation events that can occur during time required to fully encapsulate the DNA template, presuming that the nucleation events can only take place adjacent to a packaging signal. Our theory is in satisfactory agreement with the available experimental data.

**SIGNIFICANCE:** The rate and efficiency of the encapsulation of double-stranded DNA by synthetic coat proteins was recently found to be strongly enhanced by the presence of specifically positioned protein molecules on the DNA that mimic so-called packaging signals. We present a kinetic theory based on the initial stochastic nucleation and subsequent deterministic elongation of the protein coat with the aim to explain these findings. We find that equidistantly placed nucleation sites that also act as diffusion barriers on the DNA have profound and non-trivial effects, and they can either slow down or speed up encapsulation, depending on how fast nucleation is on the time scale of the elongation process. Our findings may contribute to the rational design of linear virus-like particles.

## INTRODUCTION

Considering that the filamentous or rod-like morphology is the second-most prevalent among all known viruses, we would not expect the packaging of a viral genome by coat proteins in order to produce a linear particle to be a non-trivial physics problem (2). Simply relying on the adsorption of the proteins onto the genome is not sufficient to package it effectively, not even if the adsorbed proteins attract each other, e.g., by the presence of hydrophobic patches, by hydrogen bonding or by ionic interactions mediated by multivalent ions, which in principle helps to increase the bound fraction of proteins. (3, 4) The reason is that the helical assembly of the proteins around its genome is in essence a one-dimensional process, even though the virus is a three-dimensional object. A helical arrangement or proteins in linear structures seems to be very strongly preferred, presumably in order to produce (quasi) equivalent environments to the proteins that themselves are not symmetrical objects. (5–7) Quasi one-dimensional assembly on a DNA (or RNA) template is dominated by entropy and hence prone to form “defects”, that is, naked sites on the template, which make the template vulnerable to attack by nucleases. (4, 8)

In an attempt to rationalize the successful and complete assembly of tobacco mosaic virus or TMV, probably one of the most studied viruses to date, Kraft and collaborators relied on a so-called zipper model, in which assembly can only start by the binding of a protein at a preferred position on the polynucleotide. (9) For TMV, this preferred location on the genome is called the *origin of assembly sequence* or OAS, but in the context of virology is more generally referred to as a *packaging signal* (10). The second crucial ingredient in the model is a form of allostery, where the required conformational switching of the first bound R. de Bruijn *et al.* coat protein catalyzes that of subsequently bound proteins. (6) In this case, elongation occurs by subsequent binding to already bound proteins, and the genome is packaged in a zipper-like fashion.(11)

The zipper model successfully explains how the free energy cost required to binding the first protein (or assembly of proteins) favors mixtures of completely packaged and naked templates over a mixture of partially covered templates, which would seem evolutionary beneficial (9, 11). Notably, the model does not only capture these *equilibrium* conditions, but also captures the kinetics of the whole assembly process. Based on the insights provided by this model, Hernandez-Gracia and collaborators designed a tri-block polypeptide that functions as an artificial coat protein, packages both single- and double-stranded DNA molecules with high efficiency, and protects these against breakdown by nucleases (4). Synthetic virus-like particles that are made using such designer coat proteins are already found to be promising candidates for therapeutic applications. (12, 13)

Crucial is the presence of the middle (assembly) block of the peptide that needs to undergo a conformational change from a disordered to beta sheet or beta roll configuration in order to bind to another protein, and provides allosteric control over the assembly process. (14, 15) The kinetic version of the zipper model also almost quantitatively describes the assembly kinetics of the encapsulation of the DNA by the artificial coat proteins of Hernandez-Gracia and collaborators. (11) This lends strong support for allosteric zippering as a generic mechanism for overcoming the detrimental impact of configurational entropy in quasi one-dimensional self-assembly processes involving molecular units and a template. (16)

Whilst the artificial coat protein, referred to as C-S_10_-B, does not have a specific preference for any nucleotide sequence, it commences the packaging of the DNA at one of the ends, unless the DNA is sufficiently long, in which case assembly may start at any point along the chain. (4) It seems that the ends act as (unintended) nucleation sites. Translational (or configurational) entropy can only render these ineffective if the template is sufficiently long and the entropy gain by random binding becomes appreciable.

In recent experiments, Calcines-Cruz and collaborators investigated whether introducing CRISPR-Cas12a proteins, which bind on prescribed positions on the DNA and act as barriers for the linear diffusional transport of the coat proteins along the DNA, may enhance its packaging efficiency. These proteins were also chemically linked to the aforementioned assembly block of the artificial coat protein, in which case the barriers would become *bona fide* packaging signals. (1) Remarkably, attaching both proteins to one or more positions along the DNA seem to make assembly not only more efficient but also faster, and more so if the Cas12a proteins can actually bind to the artificial coat proteins. The effect becomes stronger with increasing number of bound Cas12a proteins albeit, as it turns out, with diminishing returns.

This is actually somewhat counter-intuitive, because breaking up a long chain into (seemingly) independent shorter portions should make the assembly *less* efficient, not *more* efficient, according to the *thermodynamic* zipper model. (9, 11, 17) This, of course, calls into question the validity of the zipper model for the problem at hand. On the other hand, it would presume that thermodynamics strictly applies for the problem in hand, even though it could well be dominated by kinetics rather than thermodynamics. Indeed, it turns out that once assembled, the DNA-protein complexes and even protein-protein complexes that under appropriate conditions form in the absence of DNA, are stable against dilution. This implies that they are easier to assemble than to disassemble. (18) Incidentally, this seems to be true for icosahedral viruses too. (19) In principle, we would need to modify the kinetic zipper model of Kraft *et al.* (9), and its generalization by Punter *et al.*(11), to account for the influence of multiple “barriers” or packaging signals, and see if the model survives confrontation with the experimental data of Calcines-Cruz and collaborators (1).

Here, we opt for a simpler version that has the same basic premise, but which, in contrast to the original model, allows for analytical evaluation and relatively straightforward comparison with experiments. Our simplified model presumes a Poissonian stochastic nucleation process for the nucleation sites we define on a quasi one-dimensional template. Once binding has taken place, the model assumes elongation to occur deterministically, *i.e.*, with a fixed and constant rate. The nucleation and elongation rates are phenomenological parameters in our model, which in principle can be different for the different nucleation sites.

As our model ignores any stochasticity of the elongation process, it ignores the statistical nature of the assembly and disassembly steps during the elongation stage of the growth of the protein coat encapsulating its cargo. This we deem appropriate if the thermodynamic driving force for assembly is sufficiently strong. In addition, we do not explicitly model how precisely proteins attach to the growing end, that is, either directly from solution or by diffusion of proteins weakly bound to part of the template not yet encapsulated. (20)

In spite of these simplifications, the predictions of our simplified zipper model agree reasonably well with the experimental observations of (1). The model actually makes a number of testable predictions. Firstly, according to the model, the average protein coverage of genetic material is a monotonically increasing function of the number of packaging signals for any nucleation and elongation rate. Secondly, and perhaps counter intuitively, this results in the mean time for complete encapsulation to occur that can both decrease or increase with increasing number of packaging signals. This depends on the ratio between the elongation rate and the nucleation rate. Thirdly, we find that this mean time has an optimal value for some number of packaging signals, which is proportional the aforementioned ratio of rates.

In the remainder of this paper, we first introduce our nucleation-and-growth model for the self-assembly kinetics, and put forward a dimensionless nucleation rate that acts as the sole relevant control variable. Our model predicts the existence of three temporal regimes in the encapsulation process. For early times, template coverage scales quadratically with time. For a sufficiently low or high nucleation rate, an intermediate linear regime emerges, while for the late stages we find the template coverage to approach exponentially the state of complete coverage. Next, we introduce the barriers or assembly signals into our model, and illustrate how this influences the assembly kinetics. We show that if the dimensionless nucleation rate is sufficiently high, adding assembly signals increases the encapsulation rate. Finally, we compare our model to the experimental results of (1), and summarize our findings in the conclusion of this paper.

## METHODS

### Nucleation-and-growth model

Let each DNA strand present in the solution act as a quasi one-dimensional template of (dimensionless) length *L* that can bind *q* proteins, see Fig. 1A. We assume that the (dimensionless) length that each protein covers is small, so *m* = *L*/*q* ≪ *L*, allowing us to treat the encapsulation process in the continuum limit. Mirroring the zipper model, we assume that the nucleation of the binding of proteins onto the template occurs at one of the free ends of the DNA template. This approach is known to be valid for DNA templates that are shorter than some length that is arguably set by the binding free energy (9, 11).

**Figure 1:**
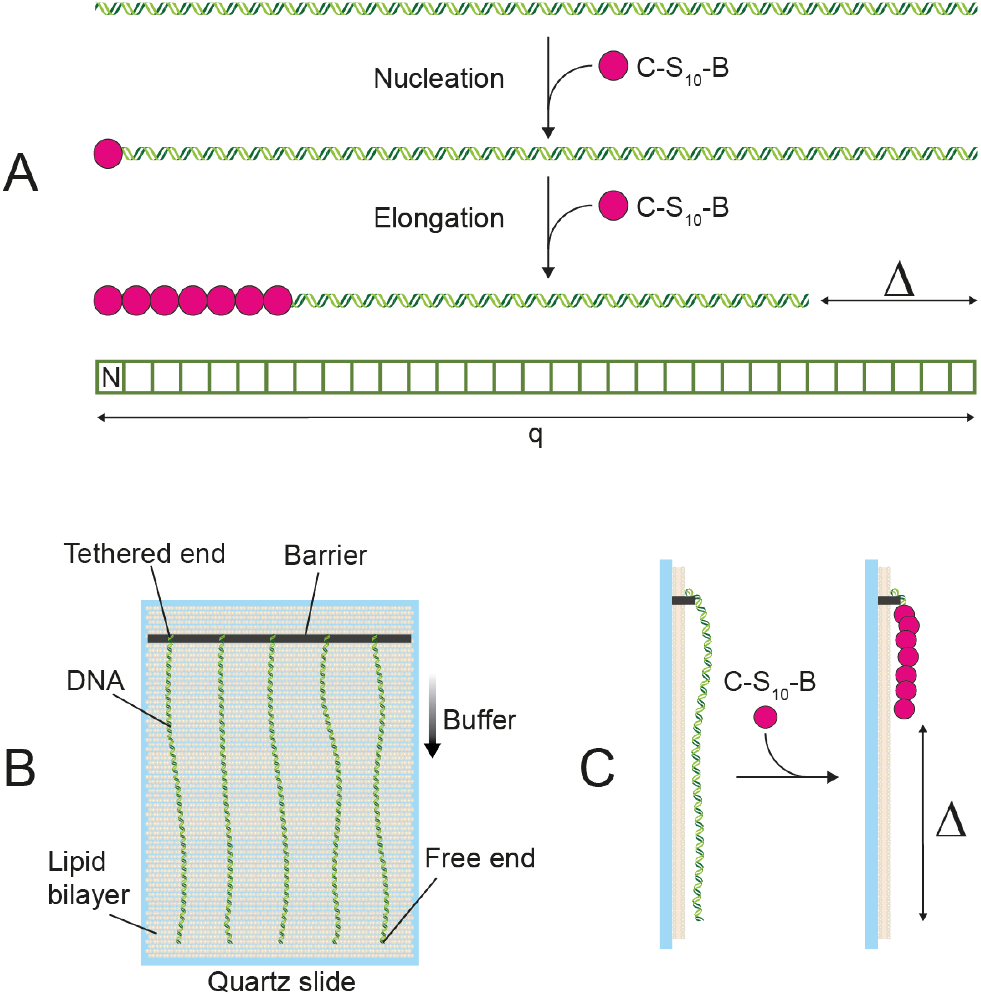
Schematic overview of our model. (A) The two-step mechanism for encapsulation of DNA by the coat protein is initiated by a stochastic nucleation process that occurs at the free end of the DNA. After this, the protein coat can engage in the elongation process, which shortens the genetic material with the length Δ by somehow folding it in the complex. The lattice representation of our model, where ‘N’ represents the nucleation site at a free DNA end and *q* is the number of binding sites on the DNA. Schematic overview of the experimental setup is shown in panels B and C. One of the DNA ends is tethered to a barrier and a buffer flow containing the C-S_10_-B coat proteins flows along the DNA chains. Coat proteins attach from this buffer solution onto the DNA templates. See Ref. (1) for more details on the experimental setup and results. Reprinted (adapted) with permission from (1). Copyright 2022 American Chemical Society.

Following established nucleation theory, we model this nucleation process as a Poisson process with a nucleation rate *I* (21). Hence, the nucleation site on the template nucleates at time *t*, that is, binds a single protein or cluster of proteins, with a probability density function given by

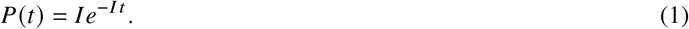

We treat the nucleation rate *I* as a phenomenological parameter that can be obtained from direct comparison of our model with experimental data, and that in principle depends on the experimental conditions such as the protein concentration (11).

After the nucleation stage, proteins attach to the nucleus and the protein coat elongates. Since the critical protein nucleus consists of a single or a few proteins, its direct influence on the fraction of the template that is encapsulated can be neglected R. de Bruijn *et al.* (20). We introduce the elongation rate *g*, presumed to be constant during the complete process. This we justify by noting that we compare our theory with experiments that are conducted at a constant protein concentration by a constant flow of protein solution over an array of DNA strands, in a setup where the fluid flow stretches the strands and the progress of their packaging can be visualized (See Fig. 1B and 1C). (1)

For simplicity, as already advertised, we treat the elongation process as if it were purely deterministic, which results in an encapsulated length *l* (*t_i_*, *t*) of the DNA template that is a linear function of time,

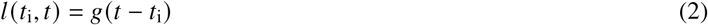

for *l*(*t_i_*, *t*) ≤ *L*. Here, *t_i_* is the time that nucleation occurs and *t* ≥ *t_i_* the time of observation. For a finite template of length *L*, the template is fully encapsulated for all *t* ≥ *t_i_* + *L*/*g*, resulting in *l*(*t_i_*, *t*) =L. We define the elongation time *t_e_* = *L*/*g* as the time required to encapsulate the whole template, if nucleation occurred at one of the free DNA ends. Experimentally, this happens to be the case as it is probably linked to the direction of the flow in the experimental setup (1).

## RESULTS

Combining the nucleation and growth stages of the encapsulation process, we define the average template coverage 〈*θ*〉 (*t*) as

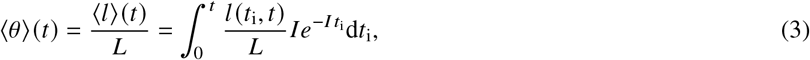

where *t_i_* < *t*. Taking the finite length of the DNA into account, this can be straightforwardly shown to yield

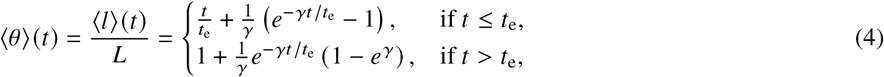

with the dimensionless nucleation rate *γ* = *It*_e_ the only relevant parameter in our nucleation-and-growth model. We note that *γ* depends on the template size via the elongation time *t*_e_ = *L*/*g*. Even though only a single nucleation event can occur per template, we can interpret γas the expected number of nucleation events that could statistically have occurred during a time equal to *t*_e_.

Whilst a seemingly simple relation, the predictions of Eq. (4) are far from that. To highlight this, Fig. 2 shows the mean template coverage as function of the scaled time *t*/*t*_e_ for selected values of the parameter *γ*. The sigmoidal shape, more conspicuous for small values of *γ*, hints at the dynamics typical of a nucleated process, which, of course, is not entirely surprising.

**Figure 2:**
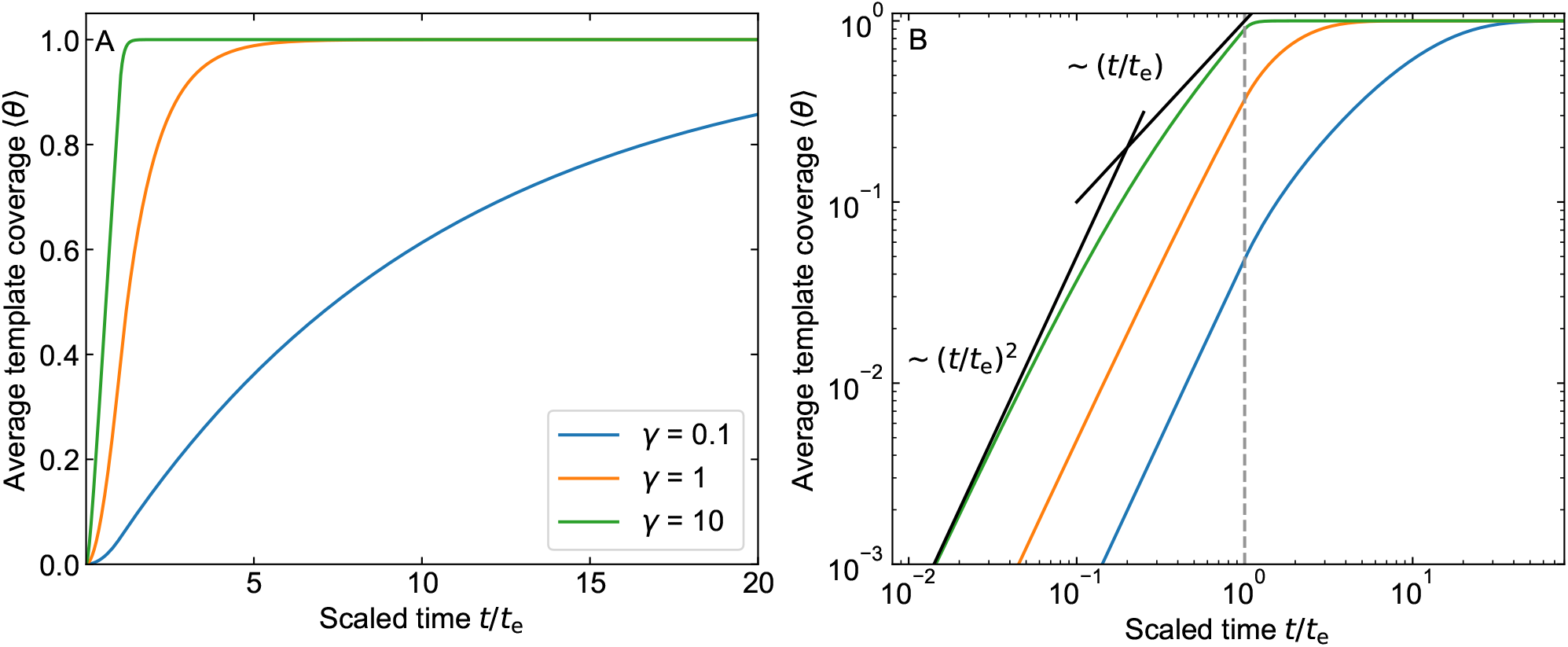
Average template coverage as function of the dimensionless time *t*/*t*_e_, with *t*_e_ the elongation time. Blue line: *γ* = 0.1, orange line: *γ* = 1, green line: *γ* = 10. Left: linear scale. Right: log scale, scaling included as guide for the eye. Grey dashed line separates initial regime (*t*/*t*_e_ ≤ 1) from exponential decay regime. Colors same as left.

From Fig. 2, we conclude that a number of distinctive growth regimes emerge and these depends on the value of *γ*. As shown in Fig. 2B, we identify an early time regime for *t* ≪ 1 ≤ *t*_e_ where the template coverage scales quadratically with time. Indeed, closer inspection of Eq. (4) shows that for this regime

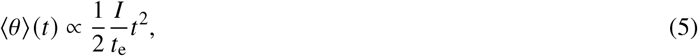

predicting that at early times encapsulation is dictated by the time scale 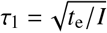.

A linear intermediate growth regime emerges if *γ* is either distinctly smaller or larger than unity, corresponding to the situation that the time scales for nucleation and growth are clearly separated. For the case that nucleation is a (relatively) fast process and *γ* ≫ 1, we find that the linear regime emerges for *It* ≫ 1, yet *t* ≤ *t*_e_. This we explain from Eq. (4) by noting that in this case the exponential term becomes small and only the linear relation survives, *i.e*.,

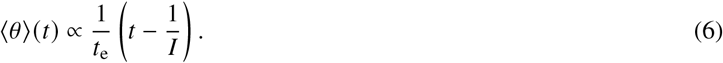

Hence, we are led to conclude that the relevant time scale must now be *τ*_2_ = *t*_e_. We interpret the 1/*I* term as a *lag time*, because elongation can only commence after the nucleation event, which on average occurs after a time 1/*I*.

For the case that nucleation is a (relatively) slow process and *γ* ≪ 1 (not shown in Fig. 2B), we find from Eq. (4) that a linear regime emerges for *t* > *t*_e_ as

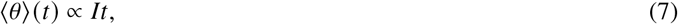

where we use that to linear order 1 - exp *γ* ~ −*γ*. The relevant time scale is *τ*_3_ = 1/*I*, as nucleation is the only relevant time scale here. This linear regime is characterized by the absence of a lag time.

Finally, for the case that *t*/*t*_e_ > 1, we find this regime to be governed by a simple exponential relaxation to complete template coverage,

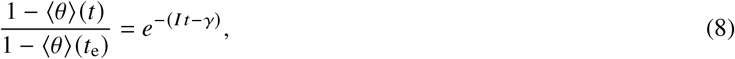

where the relevant time scale is *τ*_4_ = 1/*I*. It transpires that our seemingly simple kinetic equations produce no fewer than three regimes, governed by four time scales.

These three growth regimes actually only manifest themselves if neither the elongation nor the nucleation process fully dominates the encapsulation kinetics. If this were the case, Eq. (4) reproduces what we would naïvely expect. For the case that elongation is much faster than nucleation, if *γ* → 0, we find 〈*θ*〉 = 1 – exp(−*It*), noting that in Eq. (4) only the solution for *t* > *t*_e_ survives, that to linear order 1 – exp *γ* ~ −*γ* and that *γ*/*t*_e_ = *I*. This, in fact, is what we would expect, as it is equivalent to the (cumulative) probability that the nucleation site has nucleated before time t. For the elongation-limited case, where the nucleation process is much faster than the elongation process and *γ* → ∞, Eq. (4) reduces to *〈*θ*〉(*t*) = *t*/*t*_e_* for 0 ≤ *t* ≤ *t*_e_ and unity for *t* > *t*_e_. This, obviously, is also in accord with what we expect, for a deterministic growth process with a constant growth rate.

While the *average* template coverage provides relevant information about the encapsulation kinetics, it contains no direct information about the efficiency of *complete* encapsulation. Since genetic material is known to degrade if left unprotected, we also need to focus attention on the relevant statistics for complete template packaging. We can quantify the efficiency of complete encapsulation by considering the mean waiting time for this to happen. For any template to be completely encapsulated at time *t*, a nucleation event must have occurred *at least* a time *t*_e_ earlier. Hence, we conclude that the mean waiting time must obey

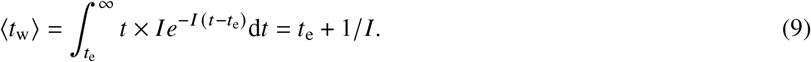

This is actually a sensible prediction and in line with our earlier analysis, if we indeed interpret 1/*I* as a *lag time* and realize that after this lag time it takes another time t_e_ to fully encapsulate the template.

Finally, since the nucleation process is stochastic, we find that the standard deviation *σ* of the completion time depends only on the quantity *I*, and obeys the simple relation *σ* = 1/*I*. From an experimental point of view, optimal control of the encapsulation process is characterized by both a short mean waiting time and a small standard deviation. Whereas the mean waiting time can be minimized by minimizing the elongation time or maximizing the nucleation rate, the standard deviation can only be made smaller by increasing the nucleation rate.

### Packaging Signals

Attaching multiple packaging signals, or protein molecules that mimic those, to the DNA template can have different effects depending on their functionality. We refer to both the inert and the chemically-active proteins as packaging signals, as both were found to enhance the encapsulation kinetics, although to a different extent. Arguably, the proteins that act as packaging signals only influence the assembly kinetics *locally*. Consequently, this suggests that these packaging signals effectively subdivide the DNA template into smaller, independent DNA templates (“sub-templates”). In principle, no distinction needs to exist between either side of a packaging signal, so growth can proceed in both directions. Bi-directional growth may cause changes in the geometry of encapsulated VLPs from linear to bend or star-like shapes, at least when multiple packaging signals (“origins of assembly”) are inserted in the RNA of TMV. (22–24) We note, however, that the experiments of Calcines-Cruz *et al*.(1) were conducted in a flow cell, in which the flow direction breaks this symmetry (See Fig. 1B). In practice, this means that the proteins move unidirectionally and that they tend to accumulate near *one* side of the packaging signal. This accumulation of weakly bound coat proteins can trigger the nucleation of a strongly bound state, akin to what happens at the free ends of the DNA. (4) The other side of the packaging signal then acts as a barrier that halts the elongation process that progresses in the direction opposite to the flow direction. (1)

To study the effect of the packaging signals on the encapsulation kinetics, we introduce in our model *n* – 1 additional, equidistantly placed packaging signals on the template. So, we have in total *n* nucleation sites, if we include the one on one of the free ends. We presume that the nucleation rates at the preferred free end of the template and those at the packaging signals are the same, which, in effect, results in all *n* portions of the template being identical. Hence, in our model the free end of the DNA acts as a packaging signal. However, as already advertised in the Introduction, the experimental findings show that specifically functionalized packaging proteins enhance the overall encapsulation rate, *i.e*., suggesting different nucleation rates for the end site and the others (1). We shall return to this issue at a later stage in this article.

Given these assumptions, we find that the average coverage of any sub-template adheres to Eq. (4), at least if we replace the template length *L* by that of the sub-templates *L/n*. Since all sub-templates are identical and independent, we conclude that the template coverage of the full template must be given by

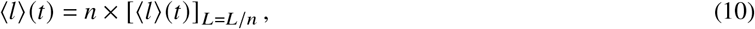

yielding

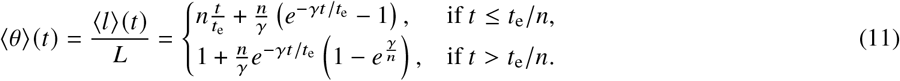

The effect of packaging signals on the average template coverage turns out to be equivalent to renormalizing both *γ* and *t*_e_ by 1/*n*, as is evident from comparing Eq. (4) and Eq. (11). This might suggest that an increase in the number of packaging signals should always produce a smaller (equal-time) template coverage. This, however, is not quite true, as we show in Fig. 3 for *γ* = 10. (Notice that the *n* = 1 case in Fig. 3 corresponds to the green curve in Fig. 2.) From the figure we conclude that increasing the number of nucleation sites n actually makes the template saturate to full coverage faster, not more slowly.

**Figure 3:**
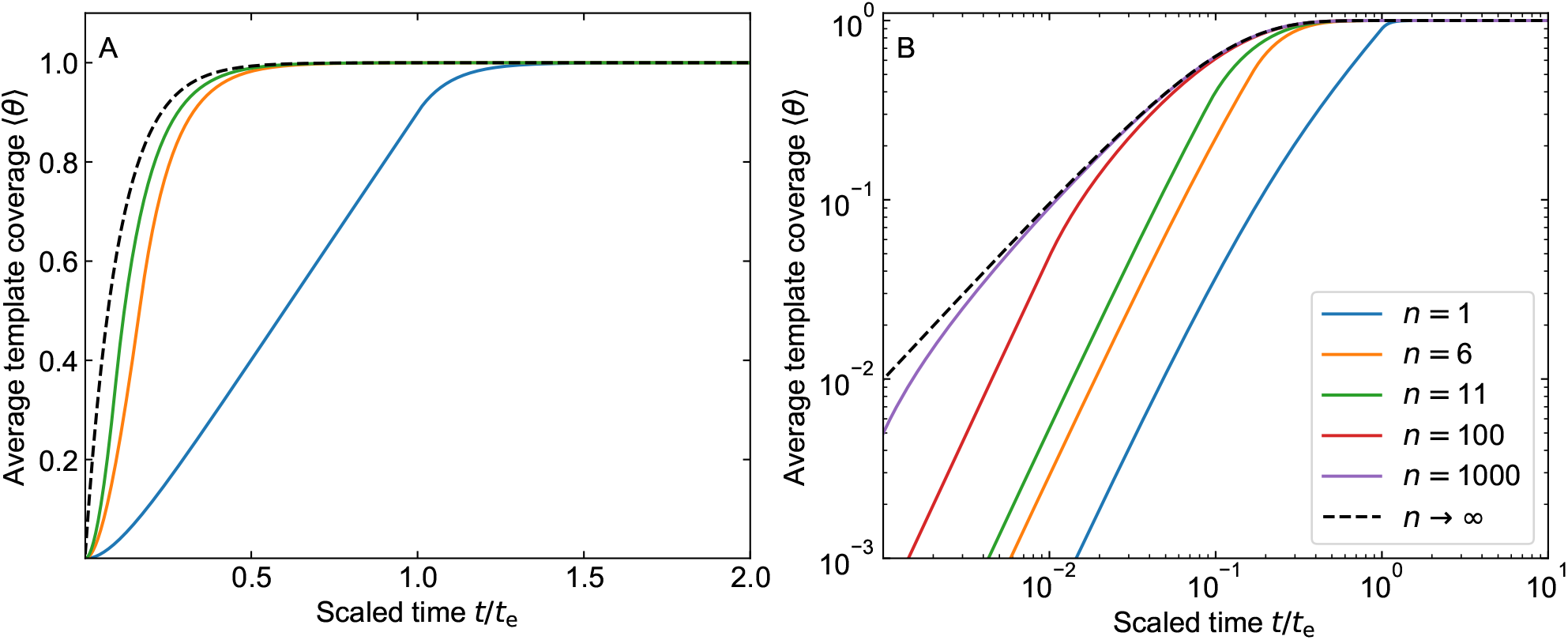
Average template coverage as function of the scaled time. Here, *t*_e_ is the elongation time, the dimensionless nucleation rate *γ* = 10 and the effect of packaging signals is included for 0 (*n* = 1), 5 (*n* = 6) and 10 (*n* = 11) and *n* → ∞ packaging signals in both graphs. Additionally, the cases for *n* = 100 (red) and *n* = 1000 are included in the right figure. Left: linear scale. Right: log scale.

Mathematically, this becomes evident if we rewrite Eq. (11) in terms of varying combinations of the time t and two time scales that emerge naturally, namely *I*^-1^and *t*_e_/*n*. The last time scale decreases with increasing *n*, arguably making the assembly faster if *t*_e_ is fixed by the length of the DNA only. A more intuitive explanation relies on realizing that it is the result of the competition between three effects. First, the *overall* nucleation rate on the template increases due to the presence of additional nucleation sites, which increases the average template coverage. Second, due to the additional nucleation sites, many different pathways now exist to achieve the same template coverage, which makes it more likely to occur. Third, the packaging signals block large parts of the template for elongation, which slows the whole process down. It turns out that the first two effects are always dominant, for Eq. (11) tells us that *∂* 〈*θ*〉 (*t*)/∂*n* ≥ 0 for all values of *γ*, *n* and *t*. We find that for large *n* there is a limit to the increase in average template coverage: for *n* → ∞ we have 〈*θ*〉(*t*) = 1 – exp(−*It*), which can be obtained from Eq. (11) in a similar fashion as we did before taking the limit *γ* → 0 for the case that *n* = 1. This limit corresponds to the case where all sites on the DNA template are encapsulated by the nucleation process only, reducing the process to a Langmuir-type adsorption process albeit without any detachment reactions, which follows a simple exponential decay, as would any first-order reaction kinetics. (20, 25)

Figs. 2 and 3 exhibit the same regimes. However, the relevant time scales do turn out to vary with the number of packaging signals *n*; for the early-time regime the relevant time scale is 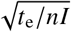 and that for the intermediate linear regime we find either *t*_e_/*n* if *γ*/*n* ≫ 1 or *I*^-1^ if *γ*/*n* ≪ 1. The time scale for the late-time exponential decay is not affected by the number of packaging signals. The *n* = 100 and *n* = 1000 curves approach the *n* → ∞ limit very slowly and a quadratic regime persists only for *t*/*t*_e_ < 1/*n*, as can be expected from Eq. (11). All of this suggests that while the whole encapsulation process appears to be faster, this is mainly true for the short- and intermediate-times.

The number of packaging signals turns out to have a significant and non-trivial impact on the time required for *complete*encapsulation. As before we measure the efficiency of complete encapsulation of the template using the mean waiting time for full coverage. The probability that the template is fully encapsulated at time t can be decomposed in the probability that *n* – 1 sub-templates have been encapsulated by time *t*, and the last sub-template *encapsulates* at time *t*. Note that for every sub-template to be nucleated at time *t*, a nucleation event must occur *at least* a time *t*_e_/*n* earlier.

Accounting for the n-fold degeneracy, which originates from the n identical pathways that now exist to reach full coverage, we find

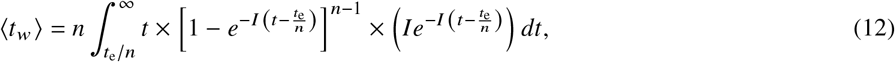

where the term between the square brackets represents the probability that certain sub-template is fully encapsulated at time *t*, and the term in round brackets denotes the probability that one of the templates encapsulates at time *t*. Carrying out the integral, we find it to reduce to

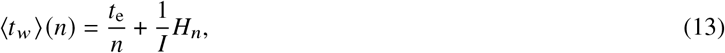

where 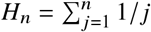 is the so-called n^th^ harmonic number. (26) We return to a discussion of the properties of *H_n_* below. We associate *I*^-1^*H_n_* with a lag time in a similar fashion as we did for the case *n* = 1 in the preceding section. It appears that this quantity also depends on the number of packaging signals on the template.

Finally, we find the standard deviation to remain inversely proportional to the nucleation rate, but now depends non-trivially on the number of packaging signals as

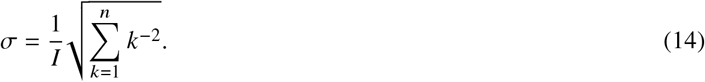

Since 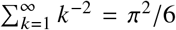 we conclude that 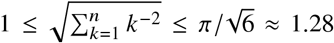 (27). While the encapsulation process should logically become more stochastic upon the addition of nucleation sites, this is not mirrored by a concomitant increase in the standard deviation, which turns out to be essentially independent of the number of packaging signals *n*.

Surprisingly, while the mean template coverage always increases with increasing number of packaging signals, this is not the case for the mean waiting time, as can be seen in Fig. 4. In the figure, we present the waiting time scaled to that of a single nucleation site, 〈*t_w_*〉(*n*)/〈*t_w_*〉(1), as a function of the number of nucleation sites *n*, for a range of values of the dimensionless nucleation rate *γ*. If this ratio attains a value smaller than unity then this indicates that the encapsulation process is more efficient, and larger than unity if it is less efficient.

**Figure 4:**
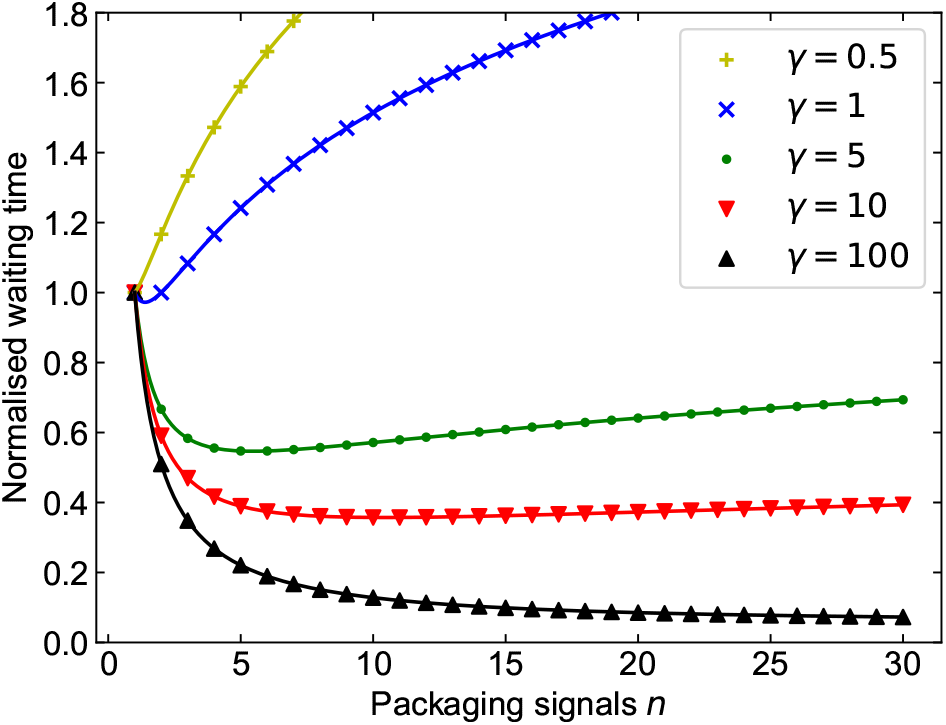
The normalized waiting time *t_w_*(*n*)/*t_w_* (1) as function of the number of packaging signals for selected values *γ* = 0.5 (yellow, plusses), *γ* = 1 (blue, crosses), *γ* = 5 (green, dots), *γ* = 10 (red, triangle-down) and *γ* = 100 (black, triangle-up). Lines are the analytic continuation of the harmonic number.

It transpires that only for *γ* > 1 the average encapsulation time decreases with increasing number of packaging signals. While this might seem inconsistent with our earlier observation that the average template coverage always increases with increasing n, this in fact is not so. To explain this, it is instructive to first focus on the case *γ* = *It*_e_ < 1, in which fewer than one nucleation event can occur in the relevant elongation time scale t_e_. Adding an additional packaging signal then effectively halts the elongation process, and the encapsulation can only continue if another nucleation event occurs. Hence, partially covered templates have a larger life time, but are easier to form as the additional packaging signals induce additional routes for partial encapsulation. Overall, this means that (on average) a larger fraction of the templates is covered, but mostly because a larger number of templates are partially encapsulated and fewer templates are completely encapsulated. Consequently, for *γ* < 1 the *complete* encapsulation is nucleation-limited.

Now, for *γ* > 1, multiple nucleation events can occur within the elongation time *t*_e_ for the complete template. Adding packaging signals can then induce nucleation before they stall the *overall* elongation process, thereby speeding up the encapsulation process. In this case, partially encapsulated templates have a shorter life time than is the case without packaging signals, and thus the mean time for complete encapsulation decreases. In other words, for *γ* > 1 the encapsulation process must be elongation-limited, not nucleation-limited.

As is evident from Fig. 4, we find that for a given dimensionless nucleation rate *γ* > 1 the encapsulation efficiency quickly decreases to an optimal value, after which it slowly increases again. We can actually determine this optimal value from Eq. (13), by using an asymptotic expansion of the harmonic number *H_n_*(26)

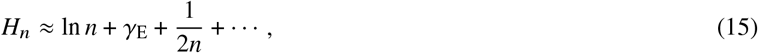

with *γ*_E_ ≈ 0.577 the Euler-Mascheroni constant. We now treat n as a continuous and as not a discrete variable, and optimize Eq. (13) to obtain

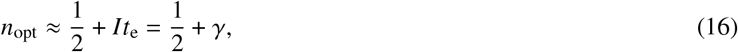

for the optimal value of the number of nucleation sites *n*_opt_.

Neglecting the constant value of 1/2, this results in an optimum encapsulation efficiency if the nucleation lag time *for a single nucleation site* is approximately equal to the elongation time of a sub-template, *i.e*., 1/*I* ≈ *t*_e_/*n*. It has to be noted, however, (i) that the decrease in the waiting time with increasing value of n is significant for the first few packaging signals, and (ii) that the optimum is very shallow, as Fig. 4 clearly shows. Indeed, for, say, *γ* = 10, any value between approximately 6 and (at least) 30 packaging signals yields a normalized waiting time that is essentially indistinguishable from its true optimum *n* =10; see Eq. (13).

From Eqs. (13), (15) and (16), we find the normalized mean waiting time corresponding to the *optimal* value of the number of nucleation sites to obey the approximate relation

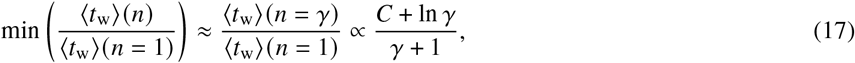

with *C* = 1 + *γ*_E_ ≈ 1.577 a constant, where we neglect a term of the order *γ*^-2^. Since the optimum in the normalized waiting time is very shallow, discreteness effects are small and treating *n* as a continuous variable does not lead to a significant error. The combination of the optimum value for *n*, and the corresponding normalized waiting time, suggests a route to optimize the encapsulation process experimentally. Obviously, this presumes that we can control the dimensionless nucleation rate *γ*. We discuss the implications of this result for the role that packaging signals may have in the assembly of naturally occurring viruses at the end of this article.

Generalizing these results to predictions of times associated with *partial* template coverage turns out to be highly non-trivial on account of the possibility of elongation processes happening simultaneously on the different sub-templates. This makes the calculation increasingly complex the larger the number of packaging signals on the template. Such a calculation would, however, still be highly interesting, as it is experimentally observable and produces information on intermediates. Actually, partial coverage times might be easier to measure if the experimentally accessible time window for measurements is, for whatever reason, too short to measure complete encapsulation.

Obviously, for most practical applications we are only interested in fully encapsulated DNA templates. Still, we can ask ourselves the question what the correlation is between the mean waiting time for partial and that for complete encapsulation. To deal with this problem, we supplement our analytical calculations with kinetic Monte Carlo (kMC) simulations, from which the mean waiting time for partial template coverage can be straightforwardly extracted. This also allows us to verify our analytical predictions, and investigate how stochasticity expresses itself in this problem.

### Kinetic Monte Carlo simulations

We set up our kMC simulations (detailed below) in such a way that they mimic the basic ingredients of our analytical theory, in which case we can simply regard our kMC results as a (near-)exact representation of our analytical model. Hence, we let the DNA template be represented by a one-dimensional lattice with a fixed number of *N*_DNA_ = 10^4^ lattice sites. This choice for *N*_DNA_ is obviously arbitrary but chosen such that it reasonably well represents the continuum approximation we use in our theoretical model. In our simulations, we introduce the nucleation reaction with a rate *k*_nuc_ and an elongation reaction with rate *k*_e_. Both reactions are stochastic, yet we do *not* allow for unbinding of proteins in the elongation process to keep as close to the model as possible. So, the overall kinetics remain based on the same model description, and the nucleation reactions can commence only on pre-designated nucleation sites that act as the packaging signals.

In our simulations, we invoke the well-known Gillespie algorithm (28–30), which can yield exact trajectories for stochastic reactions. In applying the algorithm, we calculate two random numbers at each kMC step to perform a single reaction. The first random number we use to select the reaction with a probability proportional to the reaction rate, and the second we use to select the time step for the kMC step, sampled from a Poisson distribution. These two steps we repeat until the DNA template is fully encapsulated. We refer to the review paper of Gillespie (30) and the Supplemental Material for more information (31).

We are able to map our simulations onto the parameter space of our theory by defining 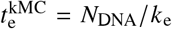 and 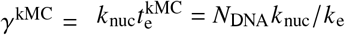. To obtain good statistics, all averages are calculated using 20,000 independent trajectories. As we show in the Supplemental Material, by setting 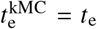 and *γ*^kMC^ = *γ*, the kMC results for the mean template coverage Fig. **S1** and mean waiting time for complete encapsulation Fig. **S2** are, for all intents and purposes, indistinguishable from the predictions of our analytical theory.

Consequently, we conclude that the stochastic nature of the elongation rate in the simulations does not appreciably influence the average observables, and that we can indeed treat the averages obtained from the kMC simulation as a (near-)exact representation of our analytical model. This can be understood intuitively if we interpret the standard deviation for a single reaction, which is inversely proportional to the reaction rate, as a measure for the stochasticity. Based on Eq. (14), we argue that the standard deviation associated with many of such reactions is essentially equal to that of a single reaction. Hence, we expect that the stochastic nature of the elongation rate becomes apparent only if *k*_el_ is of equal or larger order than *k*_nuc_, which in our case corresponds to 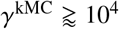.

Our kMC results for the mean waiting time 〈*t_w_*〉 as a function of the partial *targeted coverage θ* = *N*/*N*_DNA_ are summarized in Fig. 5. We present the 〈*t_w_*〉, scaled to the elongation time *t*_e_, as a function of the targeted coverage θfor the selected values of *γ*=0.1, 1 and 10. For small values of *γ*, the curves turn out to be step-like. This happens if the time between consecutive nucleation events is larger than the time required to encapsulate a sub-template. We discuss this in more detail below.

**Figure 5:**
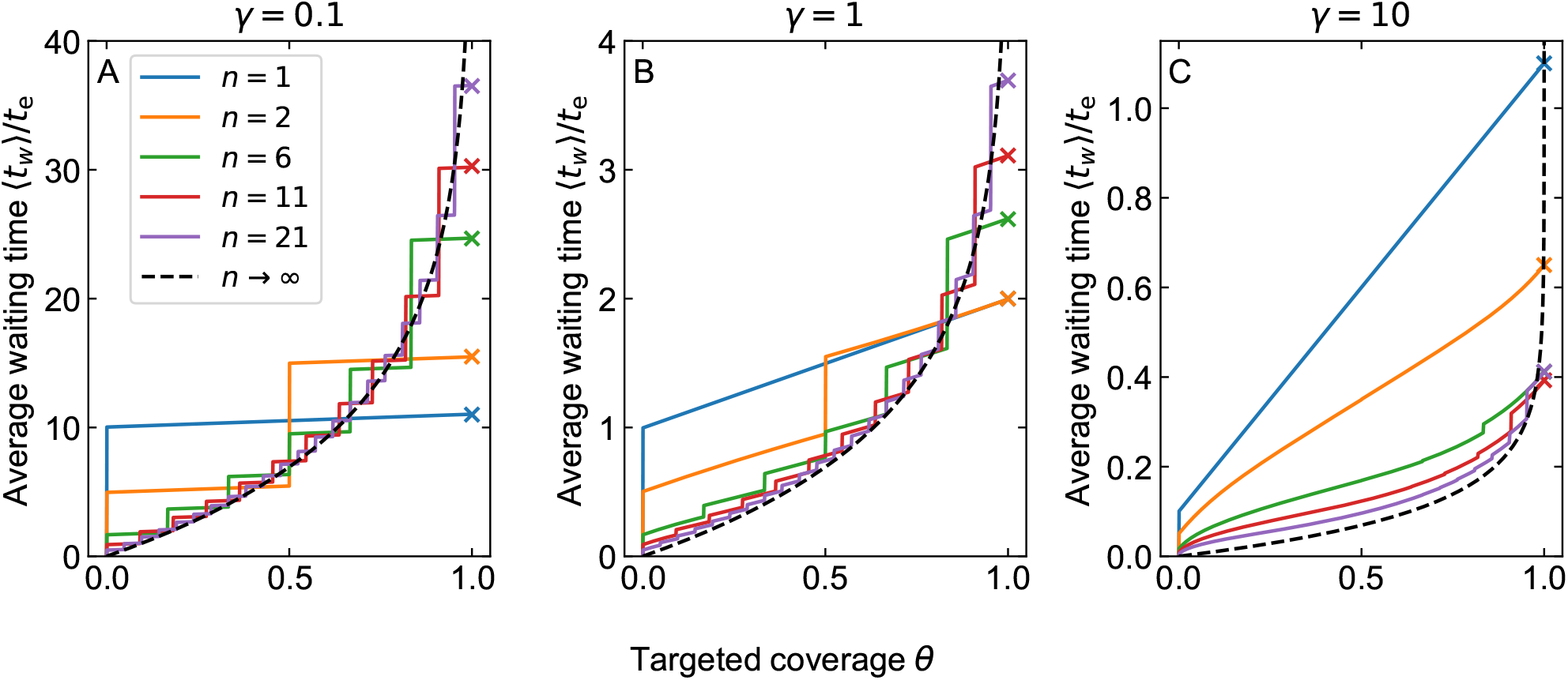
The mean waiting time to encapsulate a given fraction of the DNA template as a function of the target coverage *θ*(not to be mistaken for the average template coverage 〈*θ*〉). We include kMC results for *n* = 1 (blue), *n* = 2 (orange), *n* = 6 (green), *n* = 11 (red) and *n* = 21 (purple), and the exact asymptotic result *n* → ∞ (black, dashed). Crosses represent our analytical result and are color matched to the simulation curves. Left (A): *γ* = 0.1, middle (B): *γ* = 1 and right (C): *γ* = 10.

For the case of *n* = 1 only a single jump occurs and the mean waiting time increases linearly afterwards with a slope of unity, which holds for all finite values of *γ*. The standard deviations are not shown for the sake of clarity, but, considering that the kMC results are essentially equal to our analytical theory, we argue that the dominant contribution is from the nucleation processes only. Hence, they must be proportional to 1/*γ*.

In the figure, we have indicated our analytical prediction for complete coverage, Eq. (13), with crosses, all of which are within the 95% confidence intervals of our simulations (not shown). The prediction for the asymptotic limit for *n* → ∞, *i.e*., for the case that the template is encapsulated by the nucleation process only, is equal to 〈*t*_w_〉/*t*_e_ = −1/*γ*/*γ* log(1 − *θ*) and is shown in the figure too. (See also the Supplemental Material (31).)

The first relevant observation we make from Fig. 5, is that for small values of *γ* all curves appear to be discontinuous, that is, characterized by a sequence of steps. The reason for this is that, if nucleation events are rare, the encapsulation of a sub-template is finished before a next nucleation event happens. Only the ‘jump’ associated with the first nucleation event we expect to be infinitely sharp, because now the template changes from empty to not empty. The remaining ‘jumps’ occur as a sequence of smaller steps, which are too small to be observed in Fig. 5. In the continuum limit, this would translate to the edges of the jumps to be slightly rounded, and the slopes to be large yet remain finite. For large values of *γ*, this step-like behavior only appears to be relevant for large *n* and then only for large values of *θ*.

It turns out that we can understand the presence of these jumps if we focus attention on the mean waiting time for the nucleation events, which simplifies the description considerably. As shown in the Supplemental Material (31), the mean waiting time for the k^th^ nucleation event, given that *n* packaging signals are present on the DNA template, obeys the relation

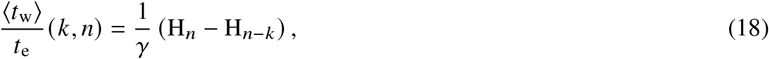

with *H_x_* again the Harmonic number. If the time between two consecutive nucleation events is sufficiently large, the encapsulation of the template may stall. This happens if the (mean) time between two consecutive nucleation events is larger than the elongation time for a sub-template, *t*_e_/*n*. Hence, we expect that a jump in the mean waiting time is present at the *k*^th^ nucleation event only if

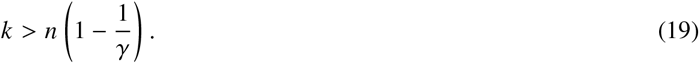

This shows that for any *γ* ≤ 1 jumps are always present, whereas for *γ* > 1, they are only present if the template coverage is sufficiently large. Comparison with Fig. 5 shows that this condition is only approximately valid for *γ* > 1, likely because several elongation processes can occur concurrently.

The second relevant observation we make from Fig. 5, is that the asymptotic limit n → ∞ appears to produce a limiting lower boundary for the average waiting time for most values of the template coverage, that is, if we exclude the presence of the jumps. As is perhaps to be expected, we find that for sufficiently small partial template coverage, adding packaging signals *always* decreases the mean waiting time, compared to the case where no packaging signals are present. Only for large template coverage does this mean waiting time increase upon adding additional nucleation sites.

We can, broadly, distinguish the crossover of these behaviors by considering the intersection between the *n* = 1 and *n* → ∞ curves, the former of which is given by 〈*t*_w_〉/*t*_e_ = 1/*γ* + *θ*, which we obtain from Eq. (9) upon replacing the encapsulation time for the complete template *t*_e_ by that of a partial template *θt*_e_. The intersection point *θ* = *N*/*N*_ONA_ is defined by 1/*γ* + *θ* = −1/*γ*log(1 – *θ*). The solution of this equation is *θ* = 1 +1/*δW*_0_(−*γ*e^−1−y^), with *W*_0_(*z*) the principle branch of the Lambert W-function.(27) From this we find that the limiting value of this intersection is given by *θ* = *N*/*N*_DNA_ = 1 – 1/e ≈ 0.63 for *γ* → 0 and by *θ* = 1 for *γ* → ∞. This suggests that adding a sufficient number of packaging signals increases the encapsulation efficiency for any value of *γ* if *θ* < 1 – 1/e, although this is not necessarily accurate for small values of *n* due to the presence of the jumps shown in Fig. 5. Surprisingly, this means that the mean time to achieve, say, one-half encapsulation *cannot* be used to quantify the effect of packaging signals on the relative efficiency to completely encapsulate the DNA template.(1)

Still, a prediction for the mean waiting time for one-half encapsulation would be interesting, if only because this observable has been determined experimentally.(1) Lacking an analytical expression that takes the number of packaging sites into account, we can make a prediction of the *shortest possible* mean waiting time to encapsulate one-half of the DNA template. In this case, we deduce from our kMC results that the shortest mean waiting time corresponds the *n* → ∞ limit. Hence, we obtain

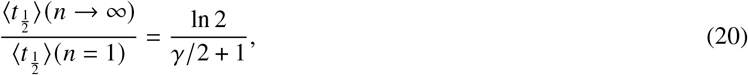

and for any finite 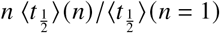 must be larger than this value. The merit of this estimate is that it depends on *γ* in a simple manner, providing us with a relatively straightforward and direct method to determine a *lower* bound on the model parameter *γ* from experiments, as we shall show at the end of the next section.

At this point, the question arises how reasonable and accurate our model is in describing actual experimental results. We answer this question by comparing our theoretical and kMC simulation results to the experiments by Calcines-Cruz *et al*.(1) in the following section.

### Comparison with experiments

Ideally, a comparison with experiments allows us not only to validate our model, but also to determine values of the model parameters *γ* and *t*_e_. As we shall see next, we find that our model agrees reasonably well with the available experimental data albeit that the values of the model parameters that we extract by curve fitting do not necessarily reflect our perhaps somewhat naive expectations. Regardless, considering the simplicity of our model, we find the agreement between theory and experiment encouraging.

As already mentioned, the experiments of Calcines-Cruz and collaborators (1) are conducted in a flow cell. This results in the DNA templates being stretched and supplied by a solution with a constant concentration of C-S_10_-B coat proteins. Proteins that attach to the templates cause these templates to reduce in length, which are believed to be fully coated when the length of the DNA strands reduces to about one-third of their initial length if no packaging signals are being used, or one-fourth if packaging signals are present on the DNA. Hence, the apparent length of the DNA can be tracked as a function of time, and can be converted to a time evolution of the fraction of the template encapsulated. For each measurement 30 minutes of observation time is taken, irrespective of whether the DNA templates are fully or only partially encapsulated in that amount of time.

We compare our model to several types of measurement by Calcines-Cruz *et al*.(1). Calcines-Cruz *et al*.(1) obtain the template coverage, averaged over 25 templates, for three cases: (i) undecorated DNA templates, *i.e*., without additional packaging signals, in contact with a C-S_10_-B coat protein solution at a concentration ranging from 10 nM to 300 nM, (ii) for a fixed C-S_10_-B concentration of 25 nM, where either 5 or 10 ‘bare’ dCas12a proteins are aimed to be attached at specific, equidistant positions on the DNA template or (iii) for a fixed C-S_10_-B concentration of 10 nM, where 5 functionalized dCas12a proteins can be attached to the DNA templates.

For the second case, the bare dCas12a-proteins are believed to act only as diffusion barriers on the DNA template, whilst in the third case, the functionalized dCas12a protein binds specifically to the silk motifs of the engineered coat proteins. We shall not attempt to curve fit the data on the latter type, as the assumption of equal binding strength must be strongly violated: the ends are likely to bind much less strongly than the packaging signals do. We return to this below.

In the remainder of this section we first compare our model with the experimental results for the mean template coverage for undecorated DNA templates. Second, we compare our model with the cases that 5 or 10 dCas12a proteins can at least theoretically be bound to the DNAs, focusing on both the mean template coverage and for the mean waiting time for one-half encapsulation.

In Fig. 6 we present the experimental data for the case without additional packaging signals for six different protein concentrations together with the confidence intervals. It should, in principle, be possible to obtain our model parameters, the elongation time t_e_ and nucleation rate *I*, by a curve fit using, *e.g*., a non-linear least-squares method (32). Unfortunately, we find this procedure to be highly sensitive to the initial parameter estimates, and does not necessarily converge unless a very good estimate of the (unknown) model parameters is supplied. Such a high sensitivity for the estimate of an initial parameter is not uncommon in non-linear least-squares methods (32).

**Figure 6:**
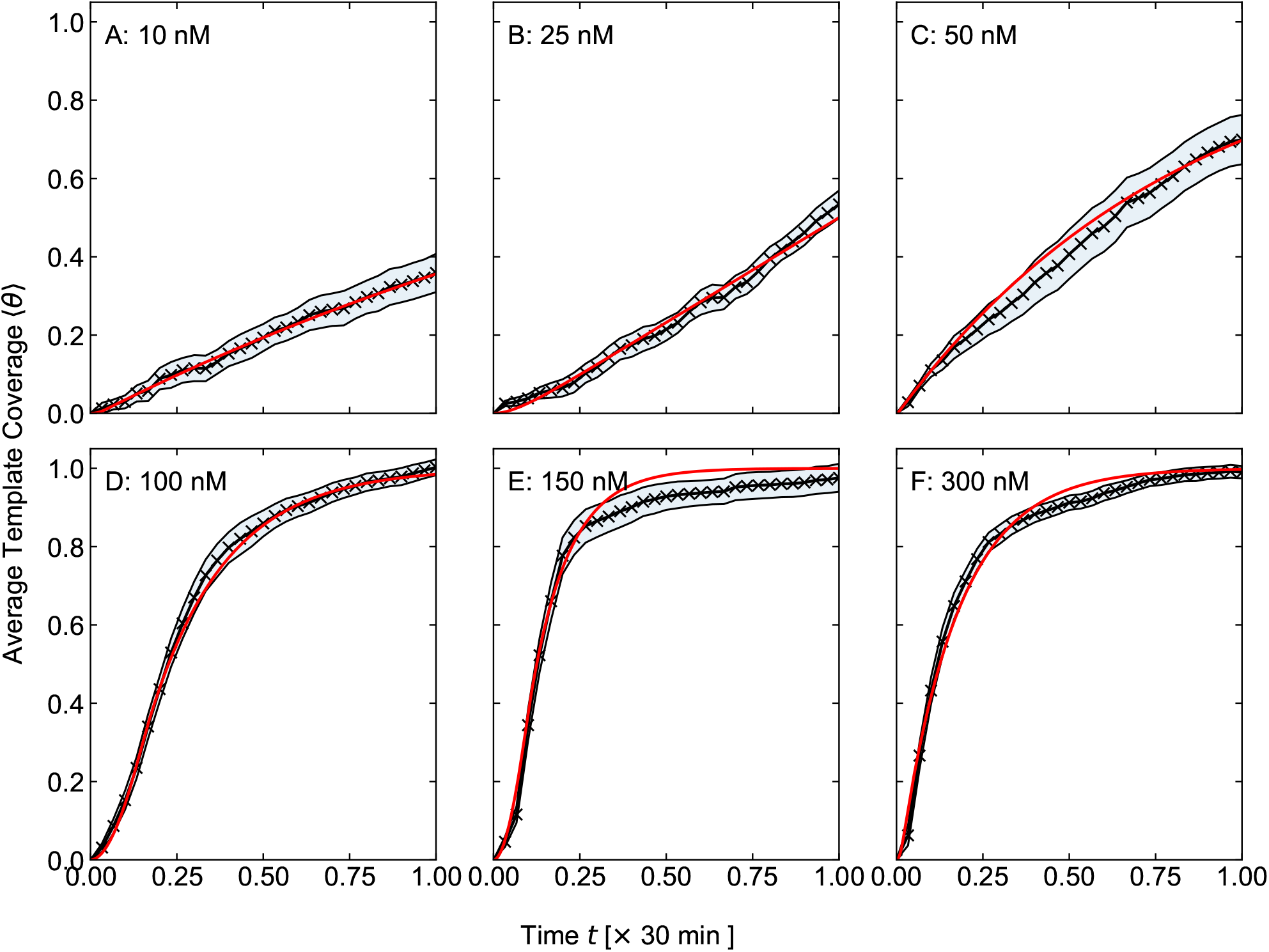
Black (crosses): experimentally obtained average template coverage 〈*θ*〉 as a function of the time t in units of 30 minutes from 25 DNA templates (1). The shaded area represents the 95 % confidence interval (1). Red (drawn line): Model fit to the average template coverage. Different curves show results for different concentrations of C-S_10_-B proteins of 10 nM (A), 25 nM (B), 50 nM (C), 100 nM (D), 150 nM (E) and 300 nM (F).

In order to work around this, we opt for a different method to find the (in some sense) best estimates for our model parameters. We first define the global residual *R*(*t*_e_, *I*) that quantifies how well our theoretical model compares with the experimental results. Since the standard errors are not constant, we use a weighted sum of squared residuals 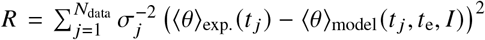, where *N*_data_ is the total number of data points, 〈*θ*〉_exp_. (*t_j_*) represents the experimental outcome at time *t_j_*, 〈*θ*〉_model_ (*t_j_*, *t*_e_, *I*) is our model value at this time and *σ_j_* is the standard error at time *t_j_*(32). We discard the first two data points that have a very small standard deviation in order to avoid that these points weigh overly heavily in our curve fitting. See SM (31) for a discussion. Next, we discretize the parameter space spanned by *t*_e_ and *I*, and determine the global residual for a set of *t*_e_ and *I* values.

We limit our region to 0 < *t*_e_ ≤ 300 [min] and 0 < *I* ≤ 33 [min^-1^], and find this to be sufficiently large. (Here, “min” stands for “minutes” in time.) Since our residual *R* is the same as the one commonly used in weighted least-squares methods, we expect that the values for *t*_e_ and *I* corresponding the smallest value of *R* are now the same as those that would have been obtained by a properly converged weighted least-squares method (32). This is obviously only true if the global minimum for R is actually within region that we investigate. Contour plots of the residuals as function of *I*(= *γ*/*t*_e_) and *t*_e_ can be found in the Supplemental Material (31).

From these contour plots we typically find that there are two types of minimum: for small values for *t*_e_ and *I* we find a relatively deep and well-defined minimum, whereas for larger values of *t*_e_ and *I* a large region with a nearly identical residual *R* is observed. The presence of these plateau-like local minima might explain why a direct curve-fit is very sensitive to the initial parameter estimates. Nevertheless, in all cases we can distinguish the global minimum for the residual *R*, and we set the model parameters *I* and *t*_e_ to the values corresponding to this smallest residual.

As shown in Fig. 6, the overall agreement between theory and experiments is good, although the theoretical prediction for 〈*θ*〉 is not always within the 95% confidence intervals for all times, in particular for the high concentrations. Our curve fits for low C-Sio-B concentrations of 10 and 25 nM are anyway less reliable and should be interpreted with some caution, because only a small fraction of the template is covered within the 30 minutes of measurement time. According to the model, the relevant time scale for early times is equal to 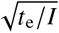, indicating that we cannot determine *t*_e_ and *I* independently if the experimental data set is limited to the early stages of the assembly process. This is also mirrored by the contour plot for the residual Fig. **S3**, where we find the residual for the global minimum to be only slightly smaller than those of the other minima. All sets of parameters including the residual *R* are presented in Table. 1.

**Table 1:** Parameter estimates for *t*_e_, *I* and *γ* = *It*_e_ for increasing coat-protein concentration C-S_10_-B if no packaging signals are attached to the DNA template. The residual *R* represents the goodness of the fit, and smaller values reflect a better fit. See main text.

| [C-S<sub>10</sub>-B] [ nM ] | $I$ [ min<sup>-1</sup> ] | $t_e$ [ min ] | $\gamma$ [ - ] | $R$ |
| --- | --- | --- | --- | --- |
| 10 | 0.015 | 1.3 | 0.02 | 4.2 |
| 25 | 0.5 | 56 | 29 | 40 |
| 50 | 0.04 | 0.2 | 0.01 | 15 |
| 100 | 0.2 | 4.1 | 0.6 | 17 |
| 150 | 0.3 | 2.7 | 0.7 | 145 |
| 300 | 0.2 | 0.6 | 0.1 | 127 |

We expect the model parameters *I* and *t*_e_ to both depend on the C-S_10_-B concentration, which is indeed what we find: see Table 1. In general, the elongation time *t*_e_ tends to decrease with increasing concentration albeit not consistently so. This global tendency of the elongation time to decrease with increasing concentration has to be expected, for a larger concentration means that the (thermodynamic) driving force for elongation must be larger. Contrasting with this, is the nucleation rate, *I*, that does not seem to vary much with increasing concentration. If we accept that the nucleation rate does indeed not depend much on the concentration at high protein concentration, this would indicate that the nucleation process is limited by some intermediate reaction-limited process rather than a diffusion-limited process (33).

Of course, we need to be cautious not to over-interpret the outcome of our curve fitting, firstly because of the simplicity of our model and secondly because the fits are based on a limited set of experimental data. And, secondly, our model ignores aspects that we know take place, such as that nucleation events at random spots on the DNA template do also occur, even though these are not included in our model.(20) Nevertheless, the overall agreement is remarkably good, indicating that our model describes most of the relevant underlying physics.

We next compare theory and experiment for the case that packaging signals have been attached to the DNA templates, which are in contact with a C-S_10_-B concentration of 25 nM. The designated binding sites are chosen such as to in theory produce 5 or 10 regularly placed on the DNA for the dCas12a-proteins to bind onto. This does not mean that all these binding sites actually do carry a dCas12a protein in the experiments (1). Experimentally, only lower bounds of 1.2 ±0.5 and 2.6 ±1.1 dCas12a-proteins could be verified to be attached to the DNA in the cases of 5 or 10 available binding sites, respectively (1). We expect actual values in between the lower bounds and the theoretical maximum.

Ignoring this inconvenience for the moment, and presuming that the theoretical number of road blocks is equal to the actual one, we show in Fig. 7 the experimental average template coverages, and the curves corresponding to the best model parameters found in our procedure. The values of these parameters we present in Table 2, noting that our model fits actually produce values for *I* and *t*_e_/*n*, which only fixes *t*_e_ if the value of *n* is known. Hence, we quote the values of *t*_e_/*n* rather than those of *t*_e_ (see also below). We further note that Fig. 7(a), representing the case for 0 dCas12a binding site, is identical to Fig. 6(b) and that we add it for the sake of reference.

**Figure 7:**
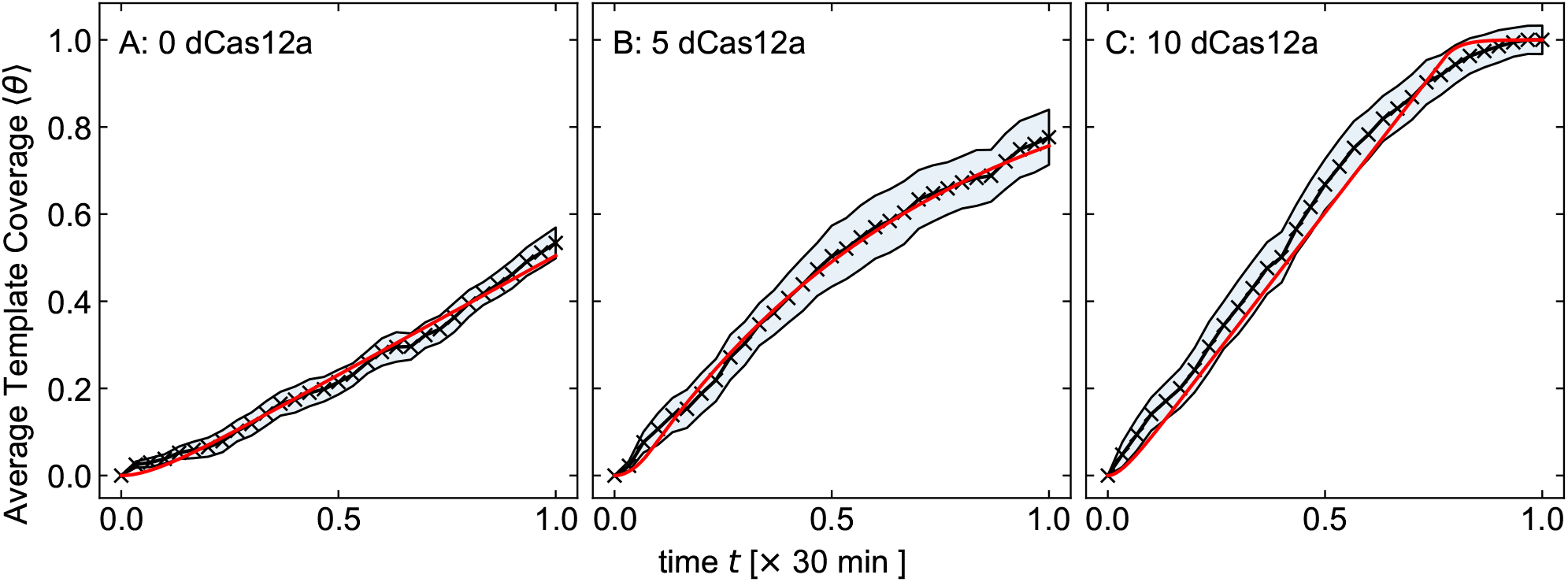
Black (crosses): experimentally obtained mean template coverage 〈*θ*〉 as function of time obtained from 25 DNA templates for [C-S_10_-B] = 25 nM (1). The shaded area represents the 95 % confidence interval. Red: Best model fit to the average template coverage. From left to right: 0, 5 and 10 dCas12a proteins attached to the DNA. For the parameter values, see Table 2.

**Table 2:** Parameter estimates for the nucleation rate *I*, the reduced elongation time *t*_e_/*n*, the elongation time *t*_e_ presuming that all dCas12a binding sites are occupied and the dimensionless nucleation rate *γ*/*n* for the case that 0, 5 or 10 dCas12a binding sites are present on the DNA template. Here, *n* denotes the number of nucleation sites on the template. The concentration of coat proteins is fixed at [C-S_10_-B] = 25 nM. The residual R represents the goodness of the fit and smaller values reflect a better fit.

| dCas12a binding sites | $I$ [$\text{min}^{-1}$] | $t_e/n$ [min] | $t_e$ [min] | $\gamma/n$ [-] | $\gamma$ [-] | R |
| --- | --- | --- | --- | --- | --- | --- |
| 0 | 0.5 | 56 | 56 | 29 | 29 | 40 |
| 5 | 0.05 | 2.5 | 15 | 0.125 | 0.7 | 6.8 |
| 10 | 1 – 30 | 24 – 26 | 264 – 286 | 24 – 720 | 264 – 7920 | 20 - 40 |

The first thing we observe is that, indeed, as announced, adding roadblocks speeds up the packaging of the DNA, and that our model is able to describe this. For 5 packaging signals we find very good agreement with our analytical model, where our curve fit is (mostly) within the 95% confidence intervals of the experiments. For 10 packaging signals agreement is somewhat less good, and we find that any nucleation rate between approximately unity and thirty per minute yields a residual *R* between 20 and 40. See also Table 2 and the Supplemental Material (31). This wide range in optimal parameter estimates implies that the fit in this particular case is not very accurate. The model curve given in Fig. 7c is that for parameter values *I* = 10 and *t*_e_/*n* = 26, which has the lowest residual *R* ≈ 20 for all values tested. Although a wide range of parameters appears to fit almost equally well, all give a value of *γ* = *It*_e_ » 1. In this case, Eq. (11) predicts that the mean template coverage 〈*θ*〉(*t*) is essentially linear until it reaches the maximum value of unity. Although the linear regime is properly captured within our model, the late-time regime clearly is not, as Fig. 7c shows.

The latter might actually not be surprising. By construction, in our model all packaging signals are equidistantly positioned on the DNA template. As already announced, it is likely that not all of the equidistant binding sites on the DNA template have a packaging signal attached to them, this assumption no longer holds. If we set the number of packaging signals in our model *n* to the *average* number of packaging signals actually present 〈*n*〉, we expect that this effect becomes pronounced only in the late-time regime. This is because the whole encapsulation process is governed by the *local* conditions only. So, for the average template coverage we can only distinguish between n uniform and n non-uniform sub-templates, if in the latter case the elongation process can encapsulate a larger part of the DNA template than would be possible in the former case. As a result of this, the effect of inhomogeneously distributed bound dCas12a molecules becomes apparent only for later times. This would explain why our model does not properly capture both short- and late-time behavior shown in Fig. 7C.

If we compare the three cases with the different number of dCas12a binding sites in Fig. 7 and Table 2, then agreement turns out to be not quite satisfactory. Indeed, within the validity of our model, we would expect that nucleation rate *I* and the elongation time *t*_e_ to not depend on *n*. From Table 2 we conclude that this is clearly not the case if we insert the theoretical values of *n* = 1,6,11, we find for *t*_e_ values of approximately 56, 15 and 250 [min], so our model appears to not be internally consistent with the data. The origins of this discrepancy may be due to (i) that the nucleation rates not being equal on the free DNA end and the packaging signals, or (ii) that the value of *n* is not equal to the maximum value and that because of that the barriers are not actually equidistant in the experiments. We return to these points at the end of the article.

Finally, we compare the experimental mean waiting time for one-half encapsulation both qualitatively and quantitatively to our model calculations. Our kMC calculations show that this mean waiting time decreases with increasing value of the number of nucleation sites *n* for *fixed* values of *γ* and *t*_e_. This is clearly in agreement with the experimental results of 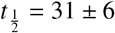 min, 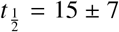 min and 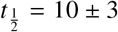 min for 0, 5 and 10 dCas12a binding sites, respectively (1). We have no direct analytical prediction how the mean one-half encapsulation time depends on the number of packaging signals, but we did conclude from Fig. 5 that the *shortest* mean waiting time for one-half encapsulation corresponded to the case where the number of packaging signals *n* becomes very large.

In that case, Equation (20) describes the relation between 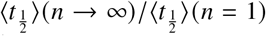 and *γ*, and can be used to derive bounds for the value of *γ*. Note that the experimental result 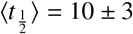 min for 10 dCas12a binding sites must now be an *upper bound* for 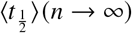. It follows from Eq. (20) that this results in a lower bound for the value of *γ*, which makes sense as a larger value for *γ* in our model correlates to a faster encapsulation process. This way we find the lower bound *γ* > 2 ± 2. Consequently, the parameter *γ* is likely to be larger than unity, and we conclude from Eq. (13) and Fig. 4 that in this particular case the packaging signals should also decrease the mean time for *complete* encapsulation.

It is important to note that this value is not completely consistent with the parameter estimations we found comparing the average template coverage. Here, we found that for the case of 5 dCas12a, *γ* = 0.7 is slightly smaller than unity, contrasting with our conclusion that *γ* is likely larger than that (See Table 2). However, since the number of packaging signals *n* is unknown, and the fit for 5 dCas12a is obtained using partial template coverage only, we refrain from over-interpreting this apparent discrepancy.

## CONCLUSION

To summarize, we have developed a nucleation-and-growth model to explain the recent experimental observation that attaching so-called packaging signals onto DNA templates can significantly increase the encapsulation rate of the DNA by designer proteins that mimic virus coat proteins.(1) These packaging signals are synthetic proteins designed with CRISPR-Cas techniques that insert themselves on predestined positions along the DNA, and can act as inert barriers or as actual binding sites for the coat proteins. In both cases, the encapsulation rate increases with the number of packaging signals, although we have limited our study to the former.

Our one-dimensional model is based on the experimental observation that the protein coat can only elongate after an initial nucleation process. Packaging signals are modeled as diffusion and elongation barriers that behave as additional nucleation sites. For the inert packaging signals, we understand this to originate from the local accumulation of coat proteins near such packaging signals in the experiments, resulting from the flow of protein solution across the stretched DNA chains. In the absence of packaging signals and for DNA molecules that are in some sense sufficiently short, nucleation happens at the far end of the DNA molecule, which then acts as an effective packaging signal.

In absence of additional packaging signals, we find that there are three growth regimes for the mean template coverage, that is, for the fraction of DNA packaged by the coat proteins. These include a quadratic early regime, a linear intermediate regime and an exponentially decaying late-time regime. This indicates that our model shows a surprisingly rich behavior considering that it is essentially determined by a single dimensionless parameter in the form of the dimensionless nucleation rate. If additional packaging signals are attached to the DNA template, we find the same three regimes but the relevant time scales for the early time and intermediate regime now depend on the number of packaging signals.

We find that the mean template coverage does always increase with increasing number of packaging signals. Interestingly, this is not mirrored by the mean waiting time for complete encapsulation, which we find to depend non-trivially on the dimensionless nucleation rate and the number of packaging signals. For sufficiently large nucleation rates, we find that this mean waiting time decreases rapidly only with the first few number of packaging signals, after which it depends only weakly on the number of packaging signals. A clear but shallow optimum exists in the mean waiting time as function of the number of packaging signals, which is important to experimentally optimize genome packaging using packaging signals.

The mean waiting time for complete encapsulation might, however, not always to be easily accessible experimentally. Our kMC simulations show, however, that the mean time for partial encapsulation is not always correlated in a simple way to the mean waiting time for complete encapsulation. In fact, based on the results presented in Figs. 4 and 5, we find that for a template coverages below approximately 63%, adding a sufficient number of packaging signals always decreases the mean waiting time with increasing number of packaging signals, even if the mean waiting time for complete encapsulation increases. Hence, there is some reason for caution interpreting experimental data on encapsulation kinetics if near completion of the process is not achieved.

The quantitative agreement between our kinetic theory and the experiments is not quite perfect. Still, we find that the *qualitative* agreement is certainly encouraging. Generally, our highly idealized model is able to explain many of the experimental results of Calcinez-Cruz and co-workers (1), suggesting that it does include most of the relevant physics. Still, that our model cannot *self-consistently* explain all experimental observations suggests that our model is incomplete. We note that some of the discrepancies might also be associated with the limited experimental data set, spanning only the first thirty minutes of the co-assembly. This becomes especially relevant for low coat protein concentrations, as only a small portion of the DNA can encapsulate within this time window. A better understanding of the disagreement between our calculations and the experiments by Calcines-Cruz *et al*. (1) would therefore also require data on the encapsulation kinetics spanning a longer time.

The first and arguably one of the main simplifications in the current model is the assumption that the nucleation sites at the free end of the DNA and at the packaging signals are equivalent, *i.e*., have the same nucleation rate. That this is not necessarily true was already shown by the experimental observation that a specifically functionalized dCas12a protein changes the overall encapsulation rate (1). This can, in principle, be incorporated relatively straightforwardly in the mean template coverage using Eq. (10) by requiring that the nucleation rate on a single sub-template differs from that on the other *n* – 1 sub-templates.

A second effect absent in our model is that not all target sites on the DNA that can bind a packaging signal, are actually bound to one. Although we argue that this should have only little influence on the early time regime, the late-time regime should be expected to be effected significantly. Indeed, it is especially in this late-time regime that our model deviates significantly from the experimental results.

The third effect that we did not incorporate into the model, is coat-protein nucleation at arbitrary places on the DNA template. As already alluded to, this has been observed experimentally for sufficiently long DNA templates(1). We expect, however, that this is mostly relevant for DNA strands without packaging signals, as the additional nucleation sites make it less likely that this process occurs. We intend to remedy all three of these simplifications in future work.

It seems sensible to remind the reader why we opted for a description based on irreversible kinetics to explain the experiments of Calcines-Cruz *et al*.(1), rather than one based on microscopic reversibility that eventually produces a thermodynamically consistent coverage. We recall that the main motivation for our model is the experimental observation that disassembly of encapsulated DNAs is exceedingly slow when exposed to protein-free buffer solution and is not observed within the experimental time (unpublished results). Actually, we also put forward a theoretical reason: adding packaging signals to the DNA template within a thermodynamic description actually decreases the *ensemble*-averaged template coverage. The implication is that from a thermodynamic point of view, the overall assembly becomes less, not more, efficient with increasing number of nucleation sites.

That this is indeed so is actually a highly non-trivial result, and due to a competition between two opposing effects. Although additional nucleation sites introduce a large entropy gain for partially encapsulated templates, the additional nucleation sites also introduce additional energy penalties for DNA encapsulation. It turns out that the energy penalty always dominates over the entropy gain. Albeit based on an equilibrium theory, we believe that lower mean coverage with increasing number of nucleation sites also implies slower dynamics, because the thermodynamic driving force for encapsulation becomes smaller. This is also in disagreement with experimental observations shown in Fig. 7.

As a final note we would like to emphasize that our work might also be relevant to understanding the role of packaging signals in naturally occurring viruses, even though we designed our model specifically to understand the effect of specifically designed packaging signals on the assembly kinetics of filamentous VLPs. Our simple model shows that adding packaging signals can speed up but also slow slow down the assembly kinetics of of *complete* viruses, and that this depends on how fast nucleation is on the time scale of elongation. See Fig. 4. This might perhaps explain the diversity in the number of packaging signals and their binding strength in viruses. It could simply be the kinetically optimal way to encapsulate the genetic material for a given interaction strength between the coat protein and the genetic material (34).

For viruses with a more complex capsid geometry, such as the icosahedral geometry, self-assembly must become a higher-than-one-dimensional process to enable the formation of complete capsids. Whether our model has any bearing on the impact of packaging signals in these viruses remains to be seen. We note, however, that in some modeling approaches the assembly process essentially reduce this from a three-dimensional to a quasi-one-dimensional process albeit with additional geometric constraints (10, 35). The complex models of Twarock and collaborators indicate that including a larger number of high-affinity packaging signals is not necessarily beneficial for the encapsulation process (35). This agrees with the results from our simple kinetic model.

All of this confirms that in molecular self-assembly, where hysteresis plays a very important role, establishing what assembled state is the most prevalent is not necessarily only dictated by thermodynamics but most certainly also by kinetics (36–38).

## AUTHOR CONTRIBUTIONS

RdB & AHG & PvdS designed the research. PCMW and RdB carried out the theoretical calculations and analyzed the data. CCC and RdB designed, conducted and analyzed the kMC calculations. TvW & PvdS analyzed the effect of packaging signals within the thermodynamic zipper model. RdB and PvdS wrote the article.

## ACKNOWLEDGMENTS

RdB. and PvdS acknowledge funding by the Institute of Complex Molecular Systems at Eindhoven University of Technology. AHG acknowledges funding from a UC MEXUS-CONACyT Collaborative grant.

## SUPPLEMENTARY MATERIAL

An online supplement to this article can be found by visiting BJ Online at http://www.biophysj.org.

