## Supplemental Material for "A nucleation-and-growth model for the packaging of genome in linear virus-like particles: impact of multiple packaging signals"

(Dated: February 23, 2022)

### I. KINETIC MONTE CARLO SIMULATIONS

Consider a one-dimensional lattice consisting of $N_{\mathrm{DNA}} = 10^4$ lattice sites. Only two types of reactions exist on this template: a nucleation reaction with rate $k_{\mathrm{nuc}}$ [$\mathrm{s}^{-1}$] and an elongation reaction with rate $k_{\mathrm{e}}$ [$\mathrm{s}^{-1}$]. We do not include a possibility for unbinding of bound coat proteins. Nucleation of coat proteins can only commence from predetermined lattice sites ("nucleation sites"). We treat the elongation process as a one-directional process. Practically, this means that at each kMC time step, the elongation reaction can only occur at the lattice site neighboring a previously encapsulated lattice site.

We have carried out our calculations using the Gillespie algorithm [1–3]. This method was initially designed to simulate stochastic (chemical) reactions for which it can, in principle, yield exact trajectories [3]. Here, we only present a short overview of this method, and we refer to Gillespie [3] for a detailed review on the subject.

Let $a_j$ represent the reaction that can occur at the j$^{\mathrm{th}}$ lattice site. At each kMC step we execute a single reaction, and associate a time $\tau$ to this kMC step. We select the reaction with a probability distribution that is proportional to the reaction rate

$$p(a_i) = \frac{a_i}{\sum_{j=1}^{N_{\mathrm{r}}} a_j}. \tag{S1}$$

Here, $N_{\mathrm{r}}$ is the total number of reactions. We select the time step by considering the probability distribution that the first reaction occurs within a time interval $\tau$. This distribution must obey the Poissonian distribution [3]

$$p(\tau) = \exp\left\{ -\left( \sum_{j=1}^{N_{\mathrm{r}}} a_j \right) \tau \right\}. \tag{S2}$$

In practice, we determine the time step and the reaction by calculating two random number $r_1$ and $r_2$ between zero and unity at each kMC time step. The first random number is used to select a reaction. We do this by finding the smallest integer $i$ that satisfies

$$\sum_{i'=1}^{i} a_{i'} > r_1 \sum_{j=1}^{N_{\mathrm{r}}} a_j, \tag{S3}$$

which can be shown to follow the proper probability distribution Eq. (S1) [3]. Using the second random number, we select the time step for the kMC step in agreement with Eq. (S2) via

$$\tau = \frac{1}{\sum_j a_j} \log\left(\frac{1}{r_2}\right). \tag{S4}$$

Since the reaction rates $a_j$ change at each kMC time step, they must be recalculated for every time step. These steps are repeated until the complete DNA template is covered by coat proteins.

The kMC calculation differs from our analytical model mainly by the elongation rate that is now stochastic. We find that this has a negligible effect on the *average* observables. To facilitate the comparison with our analytical model, we define the elongation time scale in our kMC simulation as $t_{\rm e}^{\rm kMC} = N/k_{\rm e}$, and the dimensionless nucleation rate as $\gamma^{\rm kMC} = Nk_{\rm nuc}/k_{\rm e}$, and compare for $t_{\rm e}^{\rm kMC} = t_{\rm e}$ and $\gamma^{\rm kMC} = \gamma$. To obtain reasonable statistics, we conduct 20,000 independent kMC simulations.

Our theoretical prediction for the average template coverage $\langle\theta\rangle$ agrees very well with the kMc result, as we show in Fig. S1 for the selected values for the dimensionless nucleation rate $\gamma = 0.1, 1$ and 10. In each case, we compare the cases for which there are no packaging signals ($n = 1$), 5 ($n = 6$) or 21 ($n = 20$) packaging signals. In all cases we find excellent agreement, although for $\gamma = 0.1$ the distinction between the $n = 6$ and $n = 21$ curves is small. We conclude that results of our kMC simulations and analytical theory are in excellent agreement, setting $t_{\rm e}^{\rm kMC} = t_{\rm e}$ and $\gamma^{\rm kMC} = \gamma$.

For the normalized mean waiting time $\langle t_{\rm w}\rangle(n)/\langle t_{\rm w}\rangle(n = 1)$ we find similar agreement between the theoretical and experimental results, as shown in Fig. S2. Here we limit our discussion again to the values for the dimensionless nucleation rate $\gamma = 0.1$, $\gamma = 1$ and $\gamma = 10$, and we expect similar agreement for other values. The solid lines represent the analytic continuation of Eq. (**??**) with respect to the number of packaging signals, and we include the kMC results for the selected number of packaging signals $n = 1, 2, 6, 11$ and 21. For all values shown, the kMC result is essentially indistinguishable from the analytical prediction.

### II. WAITING TIME NUCLEATION EVENTS

The step-like features of the mean waiting time as function of the template coverage that we discuss in the main text can be understood by considering the waiting time for the nucleation events only. We can obtain the mean waiting time for the $k^{\rm th}$ nucleation event by first considering the probability distribution for a state with $k$ nucleated sub-templates.

Let the DNA consist of $n$ sub-templates. Of these $n$ sub-templates, $k - 1$ *are* nucleated

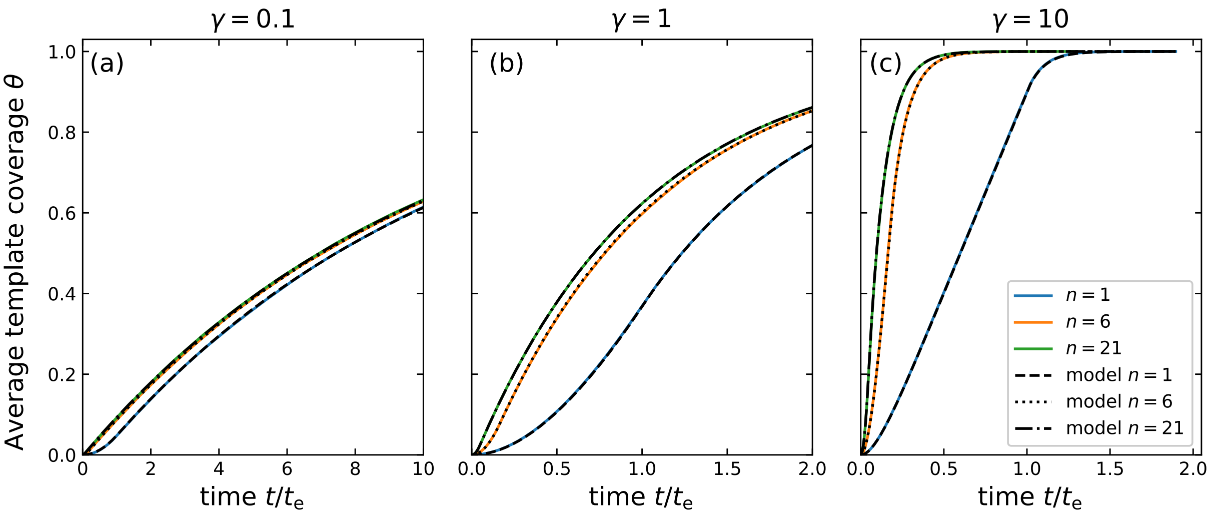


FIG. S1: Average template coverage as function of the dimensionless time $t/t_{\rm e}$, with $t_{\rm e}$ the elongation time for (a) $\gamma = 0.1$, (b) $\gamma = 1$ and (c) $\gamma = 10$. Results are shown for the case that additional packaging signals are absent ($n = 1$, in blue), and the cases where five and twenty additional ones are present. These correspond to $n = 6$, in orange, and $n = 21$, in green.

some time before time $t$ and $n - k + 1$ sub-templates are not nucleated. Since all nucleation sites are independent and identical, the degeneracy of any such state must be given by $n!/(k-1)!(n-k+1)!$. The probability distribution function for this state must therefore follow binomial statistics and is given by

$$P_{k-1}(t) = \frac{n!}{(k-1)!(n-k+1)!} \left[1 - e^{-It}\right]^{k-1} \left(e^{-It}\right)^{n-k+1} . \tag{S5}$$

Here, $1 - e^{-It}$ is the probability for a *single* nucleation site to be nucleated before time $t$ and $e^{-It}$ the probability that it is not nucleated before time $t$.

The probability density that the $k^{\rm th}$ nucleation event happens *at* time $t$ can now be found by noting that this corresponds to the case that $k-1$ templates are nucleated before time $t$, $n - k$ sub-templates are not nucleated before time $t$, and a single sub-template nucleates at time $t$. From Eq. (S5), we find this to yield

$$P_{k-1\to k}(t) = \frac{(n-k+1)n!}{(k-1)!(n-k+1)!} I e^{-It} \left[1 - e^{-It}\right]^{k-1} \left(e^{-It}\right)^{n-k} . \tag{S6}$$

The mean waiting time for the $k^{\rm th}$ nucleation event is now simply given by

$$\langle t_{\rm w} \rangle = \int_0^\infty t \times P_{k-1\to k}(t) {\rm d}t, \tag{S7}$$

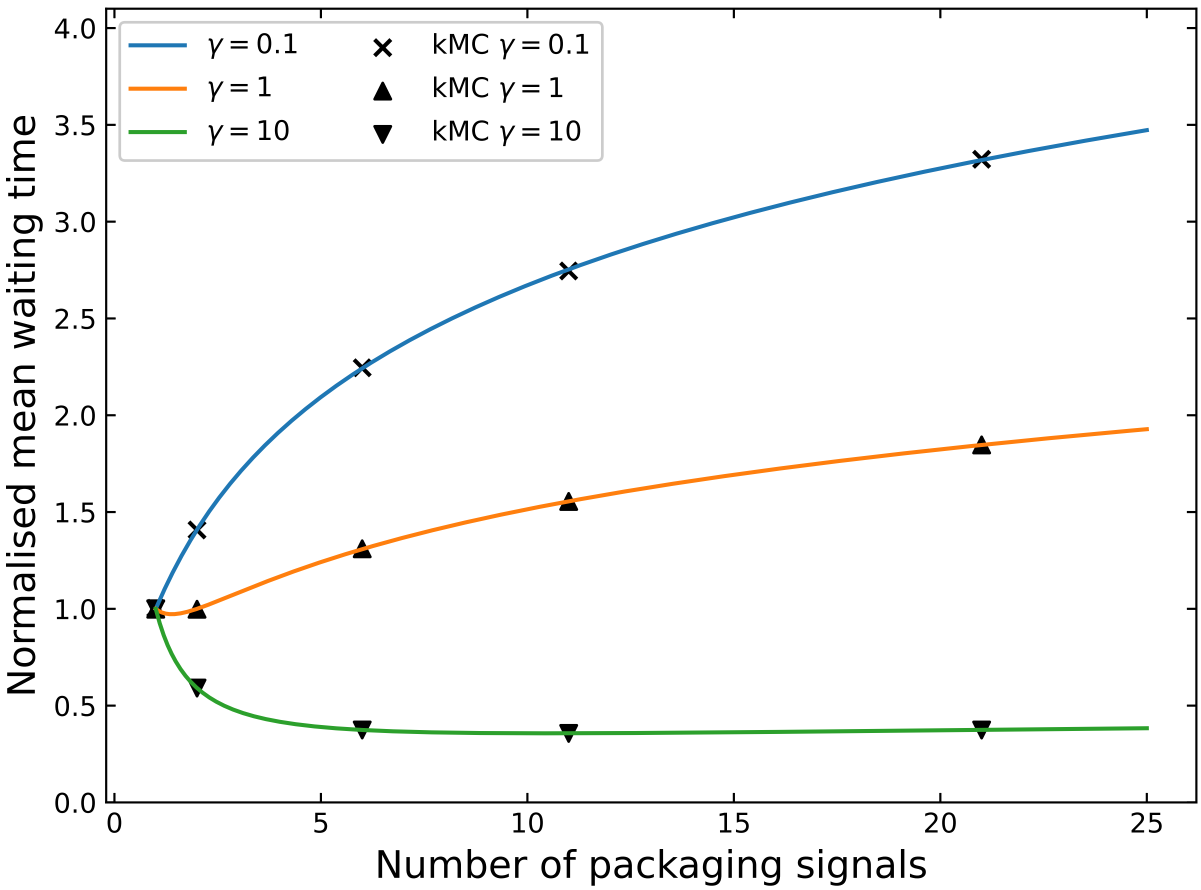


FIG. S2: Normalized mean waiting time $\langle t_{\rm w}\rangle(n)/\langle t_{\rm w}\rangle(n=1)$ as a function of the number of packaging signals $n$ on the DNA template. The lines are the analytical continuation of our theoretical prediction, whereas the markers are the results from our kMC simulations.

which yields

$$\frac{\langle t_{\rm w}\rangle}{t_{\rm e}} = \frac{1}{\gamma}\left(\mathrm{H}_n - \mathrm{H}_{n-k}\right), \tag{S8}$$

with $\mathrm{H}_n = \sum_{i=1}^{n} i^{-1}$. Using this expression, we find the (average) time difference between two consecutive nucleation events to obey

$$\Delta\left(\frac{\langle t_{\rm w}\rangle}{t_{\rm e}}\right) = \left.\frac{\langle t_{\rm w}\rangle}{t_{\rm e}}\right|_{k+1} - \left.\frac{\langle t_{\rm w}\rangle}{t_{\rm e}}\right|_{k} = \frac{1}{\gamma}\left(\mathrm{H}_{n-k} - \mathrm{H}_{n-(k-1)}\right) = \frac{1}{\gamma}\frac{1}{n-k}. \tag{S9}$$

Incidentally, the continuum limit of the mean waiting time for a DNA template where only the nucleation reaction can occur is also straightforward to determine. We first introduce the asymptotic expansion $H_x \approx \gamma_{\rm E} + \ln x$ in Eq. (S8) and next take the limit of $n \to \infty$, while keeping $k/n$ finite and constant [4]. Here, $\gamma_{\rm E} \approx 0.577$ is the Euler-Mascheroni constant.

This gives

$$\frac{\langle t_{\mathrm{w}} \rangle}{t_{\mathrm{e}}} = -\gamma^{-1} \ln k/n. \tag{S10}$$

Interestingly, the same result can also be found by simply inverting the average template coverage. Note that this only holds for the present case, where nucleation is the only process that occurs on the template. If the elongation process occurs as well, this no longer holds.

### III. FIT PARAMETER CONTOURS

Here, we present the contour plots for the comparison of our model with the experimental results. In line with the usual non-linear least-square fit procedures, we quantify the deviation between our model and the experimental results using a weighted sum of squared residuals $R = \sum_{j=1}^{N_{\mathrm{data}}} \sigma_j^{-2} \left( \langle \theta \rangle_{\mathrm{exp.}}(t_j) - \langle \theta \rangle_{\mathrm{model}}(t_j, t_{\mathrm{e}}, I) \right)^2$. Here, $N_{\mathrm{data}}$ is the total number of data points, $\langle \theta \rangle_{\mathrm{exp.}}(t_j)$ represents the experimental outcome at time $t_j$, $\langle \theta \rangle_{\mathrm{model}}(t_j, t_{\mathrm{e}}, I)$ is our model value at this time and $\sigma_j$ is the standard error at time $t_j$ [5]. Smaller values of $R$ indicate better agreement. The contour plots show the logarithm of the residual $R$ as function of the elongation time $t_{\mathrm{e}}$ and the nucleation rate $I$. Figs S3-S8 show results for increasing coat protein concentration if no dCas12a-based packaging signals are present on the DNA there is only one nucleation site at one end of the template ($n = 1$). Figs S9 and S10 showcase our results for the cases where either five or ten dCas12a binding sites are present on the DNA template, so $n = 6$ and $n = 11$ nucleation sites, for a single coat protein concentration of 25 nM.

The use of the weighted residual warrants some additional discussion. This residual clearly emphasizes data that has a relatively small error. Since the experimental data on the length of all DNA templates is normalized *per DNA chain*, the standard error is biased such that it is smaller at short times compared to late times. Note, however, that the standard errors are now both related to measurement uncertainty and to the stochastic nature of the encapsulation process itself. The latter imposes an implicit bias on the distribution of the standard errors as function of time, which we have not corrected for. This requires us to prevent that the first few data points determine the quality of the fit overly heavily, which we do by not included the first few data points. An alternative approach could be to use weights that are better suited for our particular case.

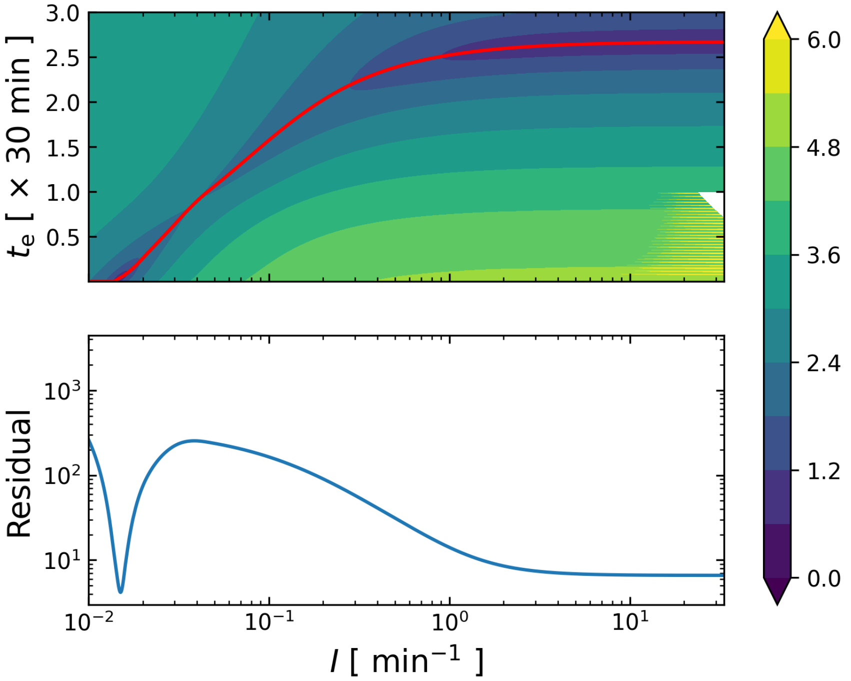


FIG. S3: Fitness landscape of our model fit to the experimental data of [6] for protein concentration [C-$S_{10}$-B] = 10 nM and a single nucleation site ($n = 1$). Top: Contour plot of the residual $R$ (see the main text) as function of the model parameters $I$ and $t_e$. Numerical values given at the color bar represent the common logarithm of the residual. Red line corresponds to the minimum residual for each $I$ value. Bottom: Residual at this red line. Two mimima present themselves for $I \approx 2 \times 10^{-2}$ and $I \gg 3$ albeit that the former represents the global minimum with the range of values of $I$ tested.

[1] D. T. Gillespie, Journal of computational physics **22**, 403 (1976).

[2] D. T. Gillespie, The journal of physical chemistry **81**, 2340 (1977).

[3] D. T. Gillespie, Annu. Rev. Phys. Chem. **58**, 35 (2007).

[4] B.-N. Guo and F. Qi, Applied Mathematics and Computation **218**, 991 (2011).

[5] T. Strutz, *Data fitting and uncertainty: A practical introduction to weighted least squares and beyond* (Vieweg+ Teubner, Wiesbaden, 2011).

[6] C. Calcines-Cruz, I. J. Finkelstein, and A. Hernandez-Garcia, Nano Letters **21**, 2752–2757 (2021).

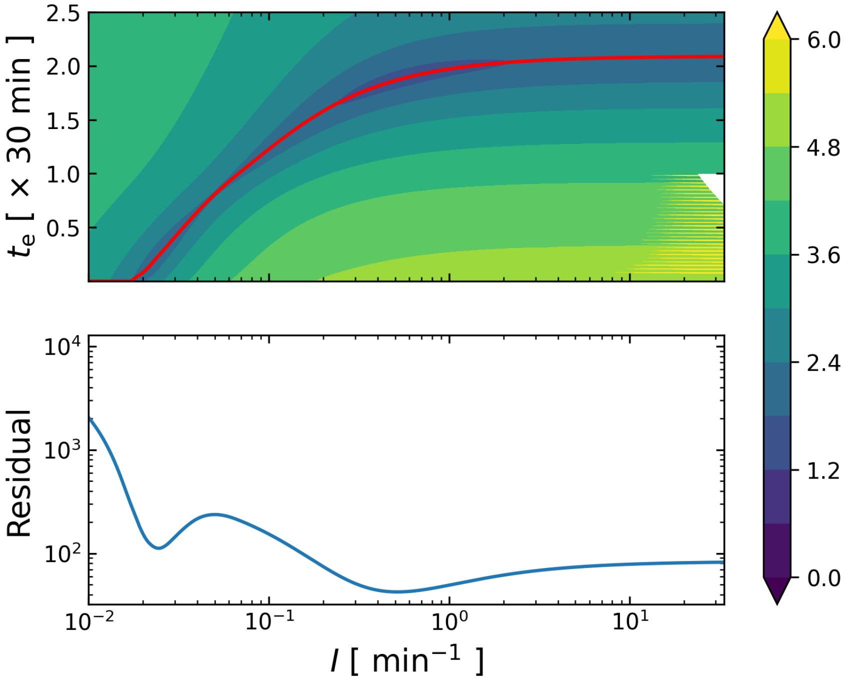


FIG. S4: Fitness landscape of our model fit to the experimental data of [6] for protein concentration [C-S$_{10}$-B] = 25 nM and a single nucleation site ($n = 1$). Top: Contour plot of the common logarithm of the residual $R$ (See main text) as function of the model parameters $I$ and $t_e$. Numerical values given at the color bar represent the common logarithm of the residual. Red line corresponds to the minimum residual for each $I$ value and the range of $t_e$ probed. Bottom: Residual at this red line. Two mimima can be distinguished, for $I \approx 2 \times 10^{-2}$ and $I \approx 5 \times 10^{-1}$, the latter representing the global minimum.

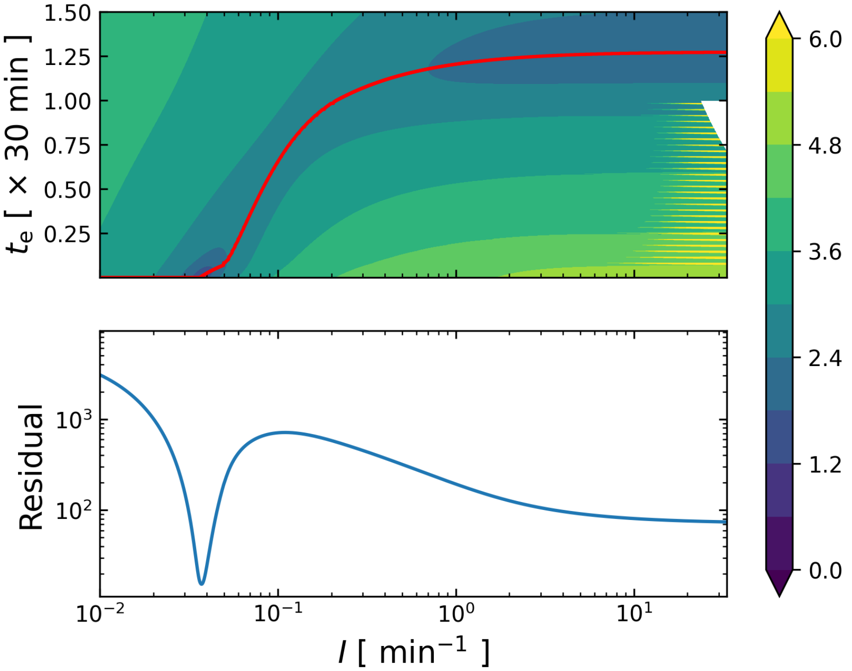


FIG. S5: Fitness landscape of our model fit to the experimental data of [6] for protein concentration [C-$S_{10}$-B] $= 50$ nM and a single nucleation site ($n = 1$). Top: Contour plot of the logarithm of the residual $R$ (see main text) as function of the model parameters $I$ and $t_e$. Numerical values given at the color bar represent the common logarithm of the residual. Red line describes the minimum residual for each $I$ value and the range of $t_e$ probed. Bottom: Residual at this red line. The global minimum is at $I \approx 4 \times 10^{-2}$, and a local minimum establishes for $I \gg 1$.

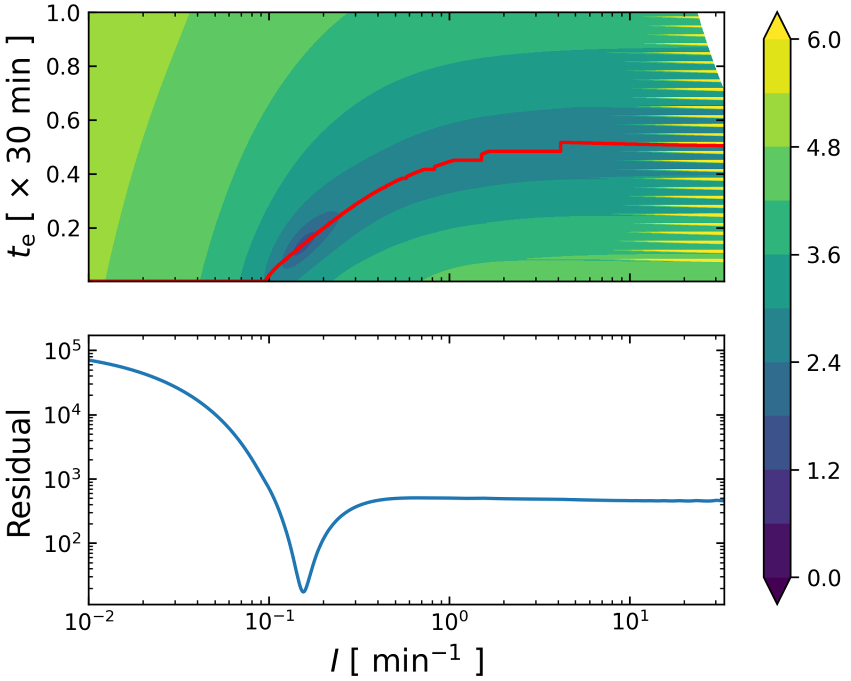


FIG. S6: Fitness landscape of our model fit to the experimental data of [6] for protein concentration [C-$S_{10}$-B] = 100 nM and a single nucleation site ($n = 1$). Top: Contour plot for the logarithm of the residual $R$ (see main text) as function of the model parameters $I$ and $t_e$. Numerical values given at the color bar represent the common logarithm of the residual. The red line describes to the minimum $t_e$ for each $I$ value and the range of $t_e$ probed. Bottom: Residual at this red line, with a clear minimum for $I \approx 2 \times 10^{-1}$.

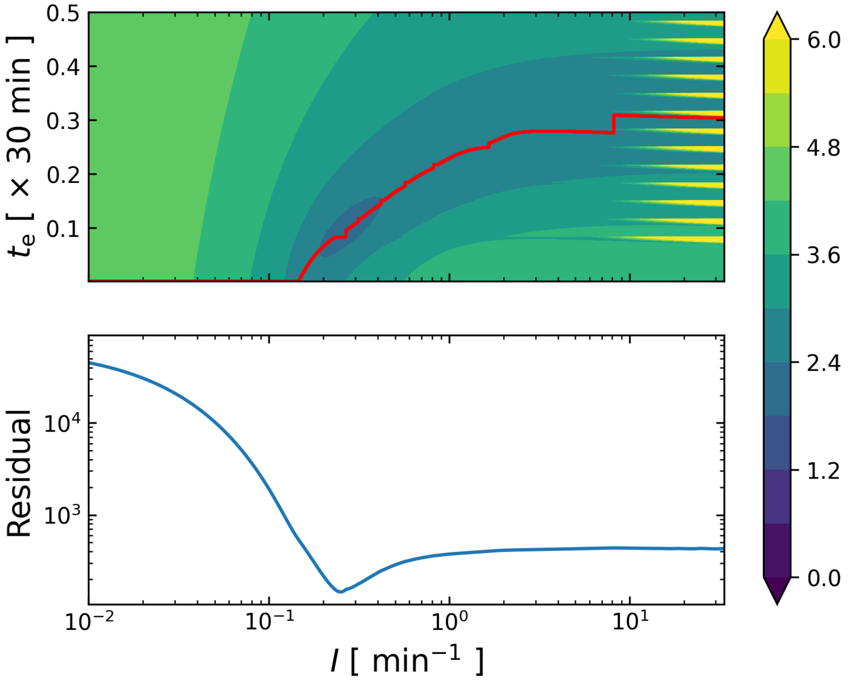


FIG. S7: Fitness landscape of our model fit to the experimental data of [6] for protein concentration [C-$S_{10}$-B] = 150 nM and a single nucleation site ($n = 1$). Top: Contour plot of the residual $R$ (see the main text) as function of the model parameters $I$ and $t_e$. Numerical values given at the color bar represent the common logarithm of the residual. Red line corresponds to the minimum residual for each $I$ value. Bottom: Residual at this red line. The global minimum is at $I \approx 3 \times 10^{-1}$.

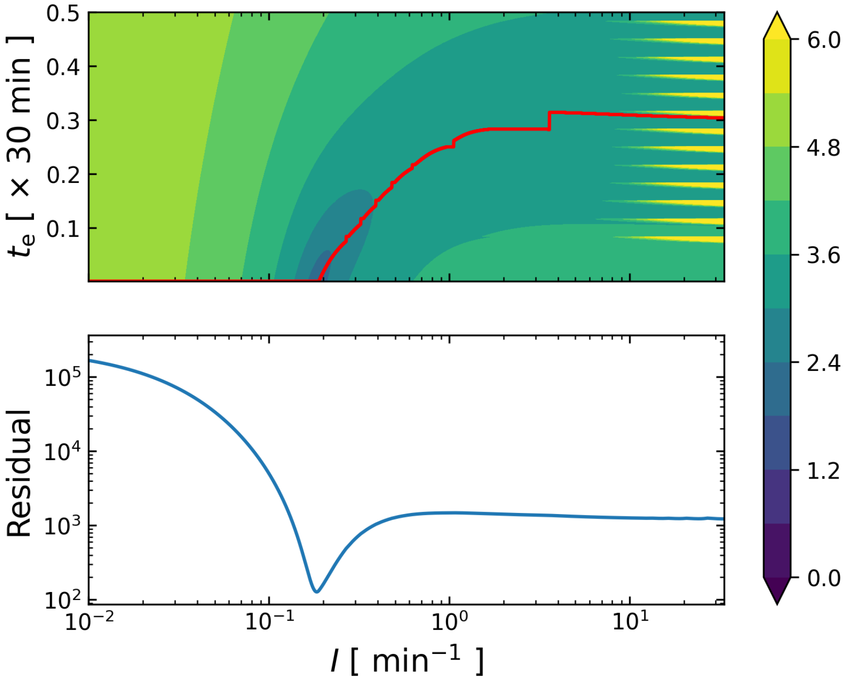


FIG. S8: Fitness landscape of our model fit to the experimental data of [6] for protein concentration [C-$S_{10}$-B] = 300 nM and a single nucleation site ($n = 1$). Top: Contour plot of the residual $R$ (see the main text) as function of the model parameters $I$ and $t_e$. Numerical values given at the color bar represent the common logarithm of the residual. Red line corresponds to the minimum residual for each $I$ value. Bottom: Residual at this red line. The global minimum is at $I \approx 2 \times 10^{-1}$.

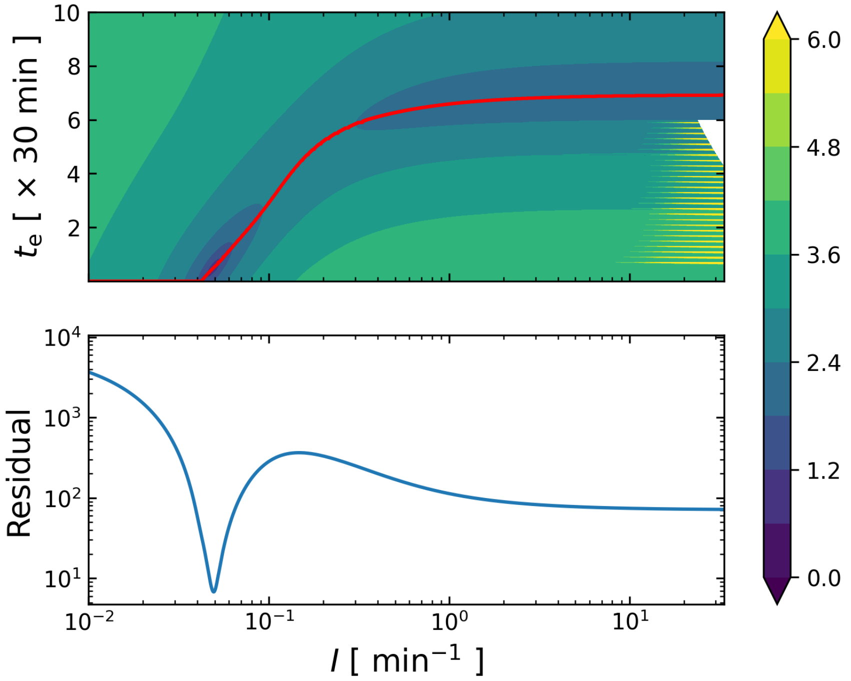


FIG. S9: Fitness landscape of our model fit to the experimental data of [6] for protein concentration [C-$S_{10}$-B] = 25 nM and five packaging signals ($n = 6$). Top: Contour plot of the residual $R$ (see the main text) as function of the model parameters $I$ and $t_e$. Numerical values given at the color bar represent the base 10 logarithm of the residual. Red line corresponds to the minimum residual for each $I$ value. Bottom: Residual at this red line. The global minimum is at $I \approx 5 \times 10^{-2}$.

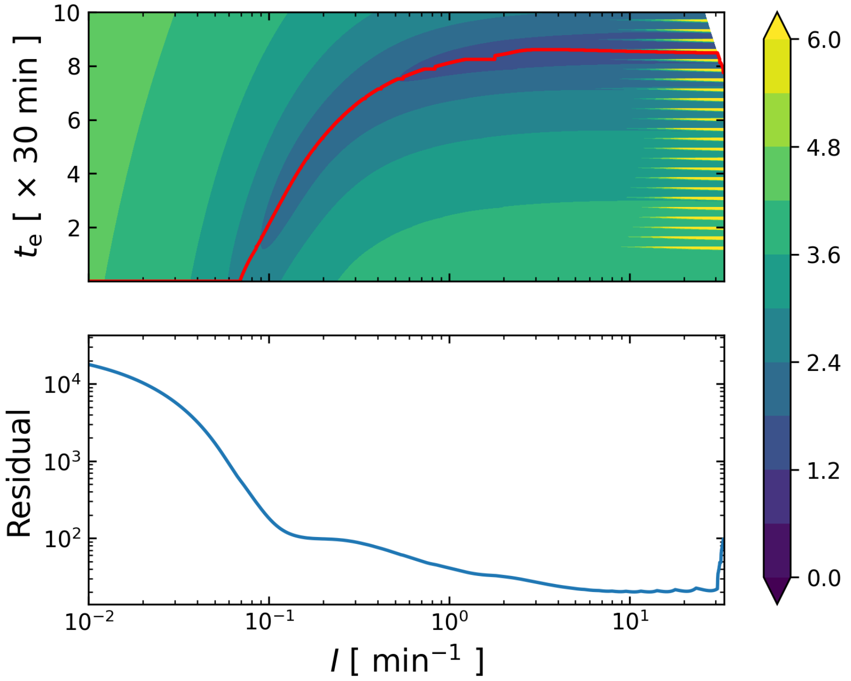


FIG. S10: Fitness landscape of our model fit to the experimental data of [6] for protein concentration [C-$S_{10}$-B] $= 25$ nM and ten packaging signals ($n = 11$). Top: Contour plot of the residual $R$ (see the main text) as function of the model parameters $I$ and $t_\mathrm{e}$. Numerical values given at the color bar represent the base 10 logarithm of the residual. Red line corresponds to the minimum residual for each $I$ value. Bottom: Residual at this red line with its minimum at $I \approx 10$.
